# NLP-12/Cholecystokinin signaling stabilizes sensory dendritic structure and protects neuronal healthspan in *Caenorhabditis elegans*

**DOI:** 10.64898/2026.03.05.709874

**Authors:** Meera M Krishna, Swapnil G Waghmare, Gabriel P Phillips, Emily C Maccoux, Tania Shaik, E Lezi

## Abstract

Aging selectively degrades neuronal structure and function, yet the signals that actively preserve neuronal integrity over adult life remain incompletely defined. In *Caenorhabditis elegans*, the PVD sensory neuron develops progressive excessive higher-order dendritic branching during normal aging that correlates with declines in proprioceptive locomotion. Using this system as a quantitative *in vivo* readout of neuronal healthspan, we identify the cholecystokinin-like neuropeptide NLP-12 as a protective signal that preserves PVD homeostasis across adulthood. *nlp-12* loss-of-function animals show early-onset excessive branching and earlier declines in proprioceptive function, whereas *nlp-12* overexpression attenuates excessive branching in aged adults without extending lifespan, indicating a neuron-focused effect on healthspan. Longitudinal analysis of an NLP-12::mKate reporter reveals age-related reduction in extracellular delivery of NLP-12 and increased retention in the soma of the DVA interneuron, where *nlp-12* is predominantly expressed. Consistent with a requirement for secretory trafficking, disrupting the NLP-12 signal peptide abolishes the rescue effects of wild-type *nlp-12* reintroduction in *nlp-12* mutants. Additionally, chemogenetic silencing of DVA during adulthood similarly accelerates excessive PVD branching, supporting an ongoing requirement for secreted DVA-derived NLP-12. Receptor genetics further show that both cholecystokinin receptor homologs, CKR-1 and CKR-2, are required downstream of NLP-12-mediated neuroprotection during aging. Finally, human cholecystokinin rescues PVD branching and partially improves proprioceptive patterning in *nlp-12* mutants, supporting functional conservation. Together, these findings define conserved cholecystokinin-like neuropeptide signaling as an adult maintenance mechanism that buffers age-associated decline in neuronal resilience.

## 1. INTRODUCTION

Neuropeptides provide an evolutionarily conserved mode of neuromodulatory signaling that can reshape circuit dynamics and behavioral output (Taghert and Nitabach 2012; van den Pol 2012). Unlike fast synaptic transmission, neuropeptides can be released from non-synaptic compartments and engage G-protein-coupled receptors (GPCRs) to influence broader cellular neighborhoods. For example, in mammalian hypothalamic oxytocin neurons, neuropeptide release can occur from dendrites and be regulated independently from axon terminal secretion, demonstrating how neuropeptide signaling can operate through extrasynaptic release and diffusion (Ludwig et al. 2002). Neuropeptide systems also play causal roles in behavioral state regulation, as shown by classic manipulations that shift feeding and sleep-wake stability when specific peptidergic pathways are perturbed (Chemelli et al. 1999; Stanley and Leibowitz 1985).

While neuropeptide signaling can influence adult neuronal function, it remains incompletely understood whether peptidergic neuromodulation broadly acts as a long-timescale maintenance program that preserves neuronal integrity as organisms age. This question is especially relevant because neuronal decline is often selective during aging. In line with this principle, distinct neuronal populations show different trajectories of vulnerability across age-related neurodegenerative diseases, suggesting that resilience could be shaped by cell-type-specific properties and by the signals neurons receive from their circuit and tissue environment (Fu et al. 2018; Kampmann 2024; Saxena and Caroni 2011). Neuropeptides are plausible contributors to such differential resilience because GPCR expression is highly cell-type specific (Garcia et al. 2015; van den Pol 2012), such that the same peptide signal can act more strongly in some neurons than others and thereby shape resilience trajectories over time.

To test whether neuron-derived peptide pathways can contribute to adult neuronal maintenance in a gene-to-phenotype framework, *Caenorhabditis elegans* (*C. elegans*) can be an ideal model, where neuromodulatory genetics can be directly linked to quantitative phenotypes *in vivo*. Importantly, genome-scale analyses identified extensive neuropeptide-like protein (*nlp*) gene families in *C. elegans*, indicating broad peptidergic signaling capacity (Nathoo et al. 2001). *C. elegans* also uses conserved neuropeptide biogenesis and processing machinery, in which proprotein convertases such as EGL-3 cleave proneuropeptides, and carboxypeptidases such as EGL-21 remove C-terminal basic residues to generate mature bioactive peptides (Jacob and Kaplan 2003).

One conserved neuropeptide family represented in this repertoire is cholecystokinin (CCK), whose receptor signaling is a canonical example of GPCR-based neuromodulation with broad physiological impact across animals. In *C. elegans*, *nlp-12* encodes a CCK-like neuropeptide that is expressed predominantly in the single interneuron DVA, a stretch-sensitive proprioceptor that modulates locomotion (Bhattacharya et al. 2014; Chen et al. 2024; Li et al. 2006). NLP-12 has been shown to signal through the GPCRs CKR-1 and CKR-2, which are homologs of mammalian cholecystokinin receptors, and prior work showed that this pathway modulates state-dependent locomotor and foraging behaviors through defined receptor-dependent mechanisms (Bhattacharya et al. 2014; Chen et al. 2024; Hu et al. 2011; Hu et al. 2015; Ramachandran et al. 2021). These ligand-receptor relationships provide a genetic framework for linking conserved neuropeptide signaling to *in vivo* adult phenotypes.

In *C. elegans*, the PVD sensory neuron provides a quantitative readout of neuronal aging. During normal aging, PVD neurons progressively develop excessive higher-order dendritic branching that correlates with age-related proprioceptive decline (Krishna et al. 2025). This phenotype reflects active remodeling rather than static overgrowth. The higher-order dendritic branches are enriched with F-actin and remain highly dynamic *in vivo*, with ongoing extension and retraction and a net bias toward excessive outgrowth in aged animals (Krishna et al. 2025). Importantly, the PVD aging phenotype can be shifted by non-cell-autonomous signals, including epidermal antimicrobial peptide signaling and extracellular matrix remodeling (E et al. 2018; Krishna et al. 2025), providing a useful context to identify additional pathways that modulate neuronal health during adulthood.

Here we show that NLP-12 functions as a protective neuromodulatory signal that promotes PVD dendrite homeostasis during adulthood. A loss-of-function mutation in *nlp-12* induces early-onset excessive higher-order dendritic branching and proprioceptive decline, while an *nlp-12* overexpression mitigates the severity of excessive branching in aged animals without affecting lifespan. Our results further indicate that secretion of NLP-12 from DVA is required for this neuroprotective effect during aging-associated remodeling. We also identify CKR-1/GPCR as a key component required for this protection, linking a defined peptide-receptor pathway to an adult-onset neuronal aging phenotype *in vivo*. Moreover, expression of human CCK can rescue the early-onset aging phenotype in *nlp-12* mutants, supporting evolutionary conservation of this pathway. Together, these findings reveal that conserved CCK-like neuropeptide signaling can support neuronal resilience during normal aging.

## 2. RESULTS

### 2.1 Loss-of-function mutation in *nlp-12* triggers early-onset aging-associated PVD excessive dendritic branching and proprioceptive deficits

The PVD neurons are a pair of bilateral mechanosensory neurons composed of an extensive dendritic arbor and a singular axon that synapses onto a few interneurons in the ventral nerve cord (Smith et al. 2010; White et al. 1986) (Fig. 1a). From the PVD soma, two primary (1°) dendritic branches extend anteriorly and posteriorly. Secondary (2°) dendrites emerge orthogonally from 1° branches at regular intervals, forming non-overlapping ‘menorah-like’ repeating units that are mainly composed of tertiary (3°) and quaternary (4°) dendrites (Fig. 1a) (Smith et al. 2010). As a polymodal neuron, PVD senses multiple stimuli, including harsh touch, proprioception, and cold temperature (Chatzigeorgiou et al. 2010; Tao et al. 2019). Previous work from our lab identified excessive dendritic branching as a robust, quantifiable hallmark of PVD neuronal aging and demonstrated that its severity is tightly correlated with aging-associated proprioceptive decline (Krishna et al. 2025). This excessive branching manifests as increasing numbers of 5° and 6° “higher-order” branches in aged animals, with a progressive increase over the course of normal aging (Krishna et al. 2025) (Fig. 1a-b).

**Figure 1:**
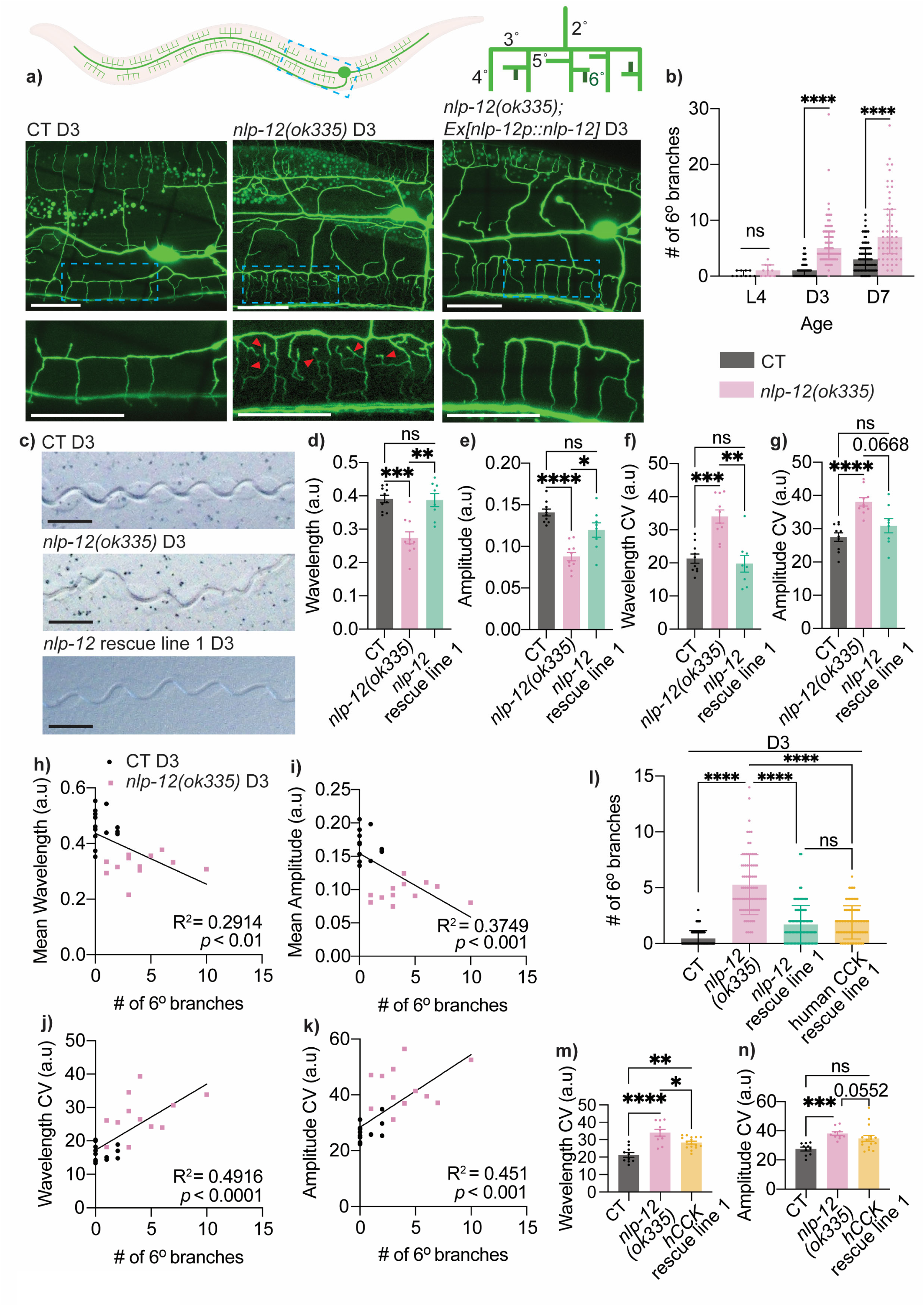
Loss of *nlp-12* triggers early-onset structural and functional aging of PVD neuron. **a)** Top left: Illustration of PVD mechanosensory neuron with a blue box indicating region imaged (created with BioRender; https://BioRender.com/xdkjwwp). Top right: Illustration of a PVD dendritic menorah unit with dark green indicating 6° dendritic branches. Bottom: Representative images of control (CT) animals [*F49H12.4::GFP(wdIs51)]*, *nlp-12(ok335)* mutants, and *nlp-12p::nlp-12* rescue in *nlp-12(ok335)* (line 1) animals at Day 3 of adulthood (D3). Red arrowheads indicate representative 6° dendritic branches. Scale bar = 25 μm. **b)** Quantification of 6° dendrites in CT and *nlp-12* mutants during aging (L4 CT: *n* = 10, L4 *nlp-*12 mutant: *n* = 10, D3 CT: *n* = 80, D3 *nlp-12* mutant: *n* = 75, D7 CT: *n* = 82, D7 *nlp-12* mutant: *n* = 53). **c)** Representative locomotion tracks from CT, *nlp-*12 mutants, and *nlp-12p::nlp-12* rescue (line 1) in *nlp-12(ok335)* animals at D3. Scale bar = 500 μm. **d-e)** Mean wavelength *(d)* and mean amplitude *(e)* measured from 100 tracks per animal and normalized to body length (CT: *n* = 10, *nlp-12* mutant: *n* = 10, *nlp-12p::nlp-12* rescue in *nlp-12(ok335)* (line 1): *n* = 8). **f-g)** Within-individual variability in wavelength *(f)* and amplitude *(g)* measured via coefficient of variation (CV) from 100 tracks per animal (CT: *n* = 10, *nlp-12* mutant: *n* = 10, *nlp-12p::nlp-12* rescue in *nlp-12(ok335)* (line 1): *n* = 8). **h-k)** Correlation between proprioception measurements and number of 6° branches at D3. Each data point corresponds to an individual animal. Black dots indicate CT, and light pink squares indicate *nlp-12(ok335)*. **l)** Quantification of 6° dendrites in CT (*n* = 76), *nlp-12* mutants (*n* = 81), *nlp-12p::nlp-12* rescue in *nlp-12(ok335)* (line 1) (*n* = 86), and human cholecystokinin (CCK) rescue in *nlp-12(ok335)* (driven by *nlp-12p* with *nlp-12* signal peptide, line 1) (*n* = 79) at D3. **m-n)** Within-individual variability in wavelength *(m)* and amplitude *(n)* measured via coefficient of variation (CV) from 100 tracks per animal (CT: *n* = 10, *nlp-12* mutant: *n* = 10, human CCK (hCCK) rescue (driven by *nlp-12p* with *nlp-12* signal sequence) in *nlp-12(ok335)* (line 1): *n* = 16) at D3. Data from additional independent transgenic lines are presented in Figure S1. Proprioception experiments performed without FUDR. Unpaired two-tailed *t*-tests with Welch’s correction were used for two group comparisons, and one-way ANOVA with Tukey’s post-hoc test or the Kruskal–Wallis test with Dunn’s correction was used for multiple-group comparisons. The Pearson’s correlation test was applied for correlation analyses. ns—not significant, * *p* <0.05, ** *p* <0.01, *** *p* <0.001, **** *p* <0.0001.

Given that this is a progressive, age-dependent phenotype, we asked whether neuromodulatory signals can act over long timescales and contribute to maintaining PVD dendritic homeostasis during adulthood. We focused on the conserved CCK-like neuropeptide NLP-12, which is produced predominantly by the proprioceptive interneuron DVA and functions in regulating locomotion behaviors (Bhattacharya et al. 2014; Chen et al. 2024; Hu et al. 2011; Hu et al. 2015; Li et al. 2006; Ramachandran et al. 2021). We found that animals carrying a loss-of-function mutation in *nlp-12*(*ok335*) exhibited early-onset PVD excessive dendritic branching at Day 3 of adulthood (D3), a young adult time point (Fig. 1a-b). Excessive branching was quantified by the number of the aforementioned 6° “higher-order” branches. Interestingly, by visual inspection, in *nlp-12* mutants, we did not detect an obvious change in age-associated dendritic beading (Fig. S1a), which is another well-established aging-related phenotype of PVD (E et al. 2018), suggesting specificity in the effects of losing *nlp-12*.

We next examined proprioceptive performance in *nlp-12* mutants and observed significant deficits at D3 (Fig. 1c-g). Proprioception, the integration of sensorimotor cues that supports posture and coordinated movement, can be assessed in *C. elegans* by quantifying the track patterns left by individual animals as they move sinusoidally across bacterial lawns (Fig. 1c). Specifically, *nlp-12* mutants showed reduced mean wavelength and amplitude of locomotor tracks (Fig. 1d-e), consistent with what has been reported by others (Tao et al. 2019). We also found further evidence of proprioceptive disruption, with *nlp-12* mutants displaying increased within-individual variability in both track wavelengths and amplitudes (Fig. 1f-g), a feature strongly associated with aging in wild-type *C. elegans* (Krishna et al. 2025). These decreased track measures along with increased within-individual variabilities resemble the gait pattern changes reported in older humans, including decreased stride length and increased step-to-step variability, which are linked to impaired proprioception (Callisaya et al. 2010; Wiesmeier et al. 2015). Consistent with our previous findings in wild-type aging animals (Krishna et al. 2025), here we also observed a strong correlation between excessive dendritic branching and proprioceptive deficits (Fig. 1h-k), supporting the functional relevance of the neuronal morphological phenotype in *nlp-12* mutants.

In order to determine whether these phenotypes reflect aging-related changes rather than developmental defects resulting from the loss of *nlp-12*, we performed a series of control experiments. While *nlp-12(ok335)* animals exhibited a shortened lifespan (Fig. S1b), we did not observe any effect on PVD structure at the L4 stage (last/4^th^ larval stage), including the number of 4° dendritic branches or the number of menorah units (Fig. S1c-d). In addition, because *nlp-12* mutants were modestly longer in body length than control animals, we normalized 6° branch numbers to individual body length and still observed significantly elevated 6° branching in *nlp-12(ok335)* animals at D3 (Fig. S1g, S1n). Moreover, after normalizing 6° branch numbers to either 4° branch numbers or menorah numbers, the excess 6° branches in *nlp-12(ok335)* animals also remained evident and followed an aging-associated progressive trajectory (Fig. S1e-f). Normalization to 4° branch numbers and menorah numbers controls for the available dendritic scaffold from which higher-order branches arise. The persistence of the phenotype under both normalizations, therefore, argues against a simple increase in arbor size and supports a selective increase in excessive higher-order branching. Given that PVD neurons are born post-embryonically and complete dendritic arborization around late L4 (Smith et al. 2010), this data supports the conclusion that the excessive dendritic branching observed in *nlp-12(ok335)* is specifically aging-associated.

Because PVD proprioceptive signaling involves mechanosensory DEG/ENaC (Degenerin/Epithelial Sodium Channel) components, we performed an exploratory analysis of *mec-10* and *del-1*, which encode DEG/ENaC subunits implicated in PVD proprioceptive function (Tao et al. 2019). At D3, *mec-10* mutants showed no obvious change in PVD 6° branch number compared to control (Fig. S1h-i). In contrast, *del-1* mutants displayed reduced 6° branches while showing similar levels of 4° branching and increased menorah units (Fig. S1h-k), indicating a structural signature distinct from the selective increase in higher-order branching observed in *nlp-12* mutants. These observations argue that *nlp-12* loss produces a branching phenotype that is not readily explained by general disruption of these mechanosensory components.

### 2.2 Human cholecystokinin (CCK) can substitute for NLP-12 to suppress excessive PVD dendritic branching in *nlp-12* mutants

We next performed rescue experiments by reintroducing wild-type *nlp-12* under its native promoter, which significantly reduced the excessive dendritic branching in mutant animals (Fig. 1a, 1l; Fig. S1o), confirming that the branching phenotype in *nlp-12(ok335)* animals is attributable to the loss of *nlp-12.* We then asked whether rescue of dendritic morphology was accompanied by improved proprioceptive performance. Re-expression of wild-type *nlp-12* also rescued all four locomotor-track measures at D3: mean wavelength, mean amplitude, and the within-individual variability of both measures were restored to levels comparable to those of age-matched control animals (Fig. 1c-g).

Thus, restoring *nlp-12* rescued both the structural and proprioceptive phenotypes of *nlp-12* mutants. NLP-12 has been proposed as a functional homolog of mammalian CCK, based on its sequence similarity to CCK-family peptides and CCK-receptor-family GPCR signaling logic in *C. elegans* (Bhattacharya et al. 2014). To directly test its functional conservation in our PVD aging phenotype, we asked whether human CCK could substitute for NLP-12 *in vivo*. We expressed human CCK cDNA under the *nlp-12* promoter together with the *nlp-12* signal sequence to support proper secretion. This human CCK expression significantly reduced excessive dendritic branching in *nlp-12(ok335)*, to a similar extent as *C. elegans nlp-12* rescue (Fig. 1l; Fig. S1p). We next asked whether this structural rescue extended to proprioceptive performance. Our data showed that, while mean wavelength and mean amplitude in human CCK rescue animals remained similar to those of *nlp-12* mutants, there was a partial rescue of wavelength coefficient of variation and a similar trend for amplitude coefficient of variation (Fig. 1m-n; Fig. S1l-m). Thus, human CCK rescued PVD morphology and recapitulated selected aspects of the proprioceptive effects of native NLP-12. Compared with the broader rescue produced by wild-type *nlp-12*, the incomplete functional rescue by human CCK may reflect differences in the strength, temporal pattern, processing, or circuit distribution of CCK-like signaling. Taken together, our findings support evolutionary conservation of CCK-like neuropeptide signaling in maintaining neuronal structural integrity and modulating proprioceptive performance during aging.

### 2.3 Aging is associated with reduced NLP-12 secretion, which is required for NLP-12 regulation of PVD dendritic branching

While we did not observe obvious aging-related alterations in *nlp-12* expression in wild-type animals (Fig. S2a), neuropeptide signaling depends not only on production but also on appropriate secretion. We therefore asked whether NLP-12 secretion from DVA neurons changes with age and whether secretion is required for NLP-12-mediated protection against aging-related changes in PVD dendritic integrity. To monitor NLP-12 secretion *in vivo*, we first constructed a translational reporter expressing NLP-12::mKate under the *nlp-12* promoter. We simultaneously labeled coelomocytes, which are six scavenger cells in *C. elegans* positioned near the head, mid-body, and tail that continuously endocytose pseudocoelomic fluid and internalize secreted proteins (Fares and Greenwald 2001) (Fig. 2a). Leveraging this system, we examined NLP-12::mKate fluorescence in coelomocytes as a proxy for secreted NLP-12 levels, while the fluorescence signal associated with the DVA soma region provided an estimate of intracellular retention. To verify the identity of the soma-associated signal, we crossed the NLP-12::mKate reporter strain with ZM11177 *[twk-40p(short)::eGFP + myo-2p::wCherry]*, which expresses cytosolic GFP in DVA. In the resulting animals, the soma-associated mKate signal was contained within the GFP-defined boundaries of the DVA cell body (Fig. S2b). We next used this reporter to examine whether NLP-12 secretion changes with age. To control for inter-individual variability, we used a longitudinal imaging design in which each animal was imaged at D3, recovered and maintained separately, and then reimaged at D7. With this within-animal comparison, we observed reduced NLP-12::mKate accumulation in coelomocytes with age, accompanied by increased signal at the DVA soma region (Fig. 2b-d; Fig. S2c), suggesting an age-associated reduction in NLP-12 secretion.

**Figure 2:**
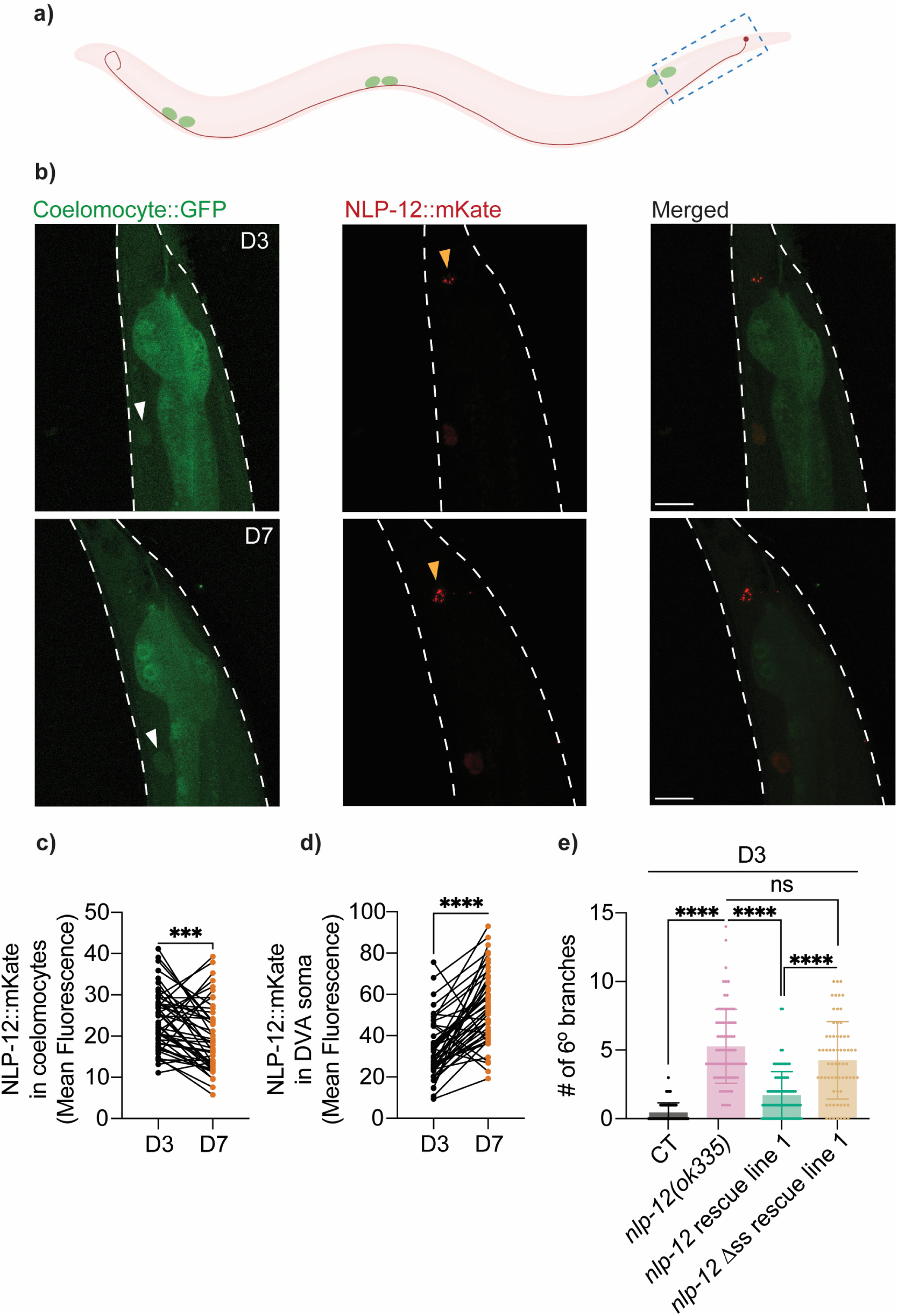
Secretion of NLP-12 is critical for its role in regulating PVD dendritic branching. **a)** Illustration of DVA neuron and coelomocytes. Three pairs of coelomocytes indicated in green, and DVA soma and neurite indicated in red. Blue dashed box indicates region shown in representative confocal images in *(b)*. **b)** Representative images of coelomocytes and DVA interneuron in D3 and D7 CT animals *[unc-122p::GFP, nlp-12p::nlp-12::mKate]*. *unc-122p::GFP* was used to visualize coelomocytes. White arrowheads indicate coelomocytes. Yellow arrowheads indicate DVA soma location. White dashed lines indicate the outline of an animal. Mild gut autofluorescence, a normal physiological feature in *C. elegans*, appears throughout the body. Scale bar = 25 μm. **c-d)** Quantification of mean gray value of NLP-12::mKate fluorescence averaged across all coelomocytes *(c)* and within DVA soma *(d)* at D3 and D7 (*n =* 48 individual animals measured longitudinally at both ages; each connecting line represents paired measurements from the same individual animal). **e)** Quantification of 6° dendrites in CT (*n* = 76), *nlp-12(ok335)* (*n* = 81), *nlp-12p::nlp-12* rescue in *nlp-12(ok335)* (line 1) (*n* = 86), and *nlp-12p::nlp-*12 *Δss* (mutated signal sequence) rescue in *nlp-12(ok335)* (line 1) (*n* = 66) at D3. Data from additional independent transgenic lines are presented in Figure S2. The Wilcoxon matched-pairs signed rank test was used for paired comparison, and the Kruskal–Wallis test with Dunn’s correction was used for multiple group comparison. ns—not significant, *** *p* <0.001, **** *p* <0.0001.

To test the importance of secretion in NLP-12’s function in limiting PVD excessive branching, we engineered a secretion-defective *nlp-12* rescue construct by mutating the signal peptide, substituting the first four amino acids of the hydrophobic core with hydrophilic glutamic acids. Disrupting the hydrophobic core is expected to impair signal peptide function and thereby reduce entry of NLP-12 into the secretory pathway. To assess the effect of this mutation on protein localization, we examined corresponding mKate-tagged reporters. All animals expressing wild-type NLP-12::mKate showed a detectable signal in both the DVA soma and coelomocytes; in contrast, all NLP-12Δss::mKate (signal-peptide mutant) animals retained a signal in the DVA soma, but none showed a detectable signal in coelomocytes (Fig. S2d). We next tested whether the signal-peptide mutation also impaired *nlp-12* function, and found that expression of the NLP-12Δss::mKate construct failed to reduce the excessive PVD dendritic branching phenotype of *nlp-12(ok335)* animals, whereas wild-type NLP-12::mKate successfully rescued the branching phenotype (Fig. 2e; Fig. S2e). Together, these findings indicate that NLP-12 secretion is required for its protective effect on PVD dendritic structure and suggest that an aging-associated reduction in NLP-12 secretion may contribute to the aging-associated excessive dendritic branching of the PVD neuron.

### 2.4 Overexpression of *nlp-12* or human CCK attenuates aging-associated PVD excessive dendritic branching

Given the age-associated reduction in NLP-12 secretion, we next asked whether elevating *nlp-12* is sufficient to counteract aging-associated PVD excessive branching. Transgenic overexpression of *nlp-12* driven by its native promoter in wild-type animals did not alter the 6° branch number at D3 but significantly reduced excessive dendritic branching at D7; this protective effect was also observed at D9 (Fig. 3a-d; Fig. S3a). This temporal pattern suggests that *nlp-12* overexpression attenuates age-associated dendritic remodeling rather than altering the young-adult PVD arbor development. Interestingly, *nlp-12* overexpression did not alter lifespan (Fig. 3e; Fig. S3b), implying that its protection against neuronal structural aging is separable from broad effects on organismal longevity.

**Figure 3:**
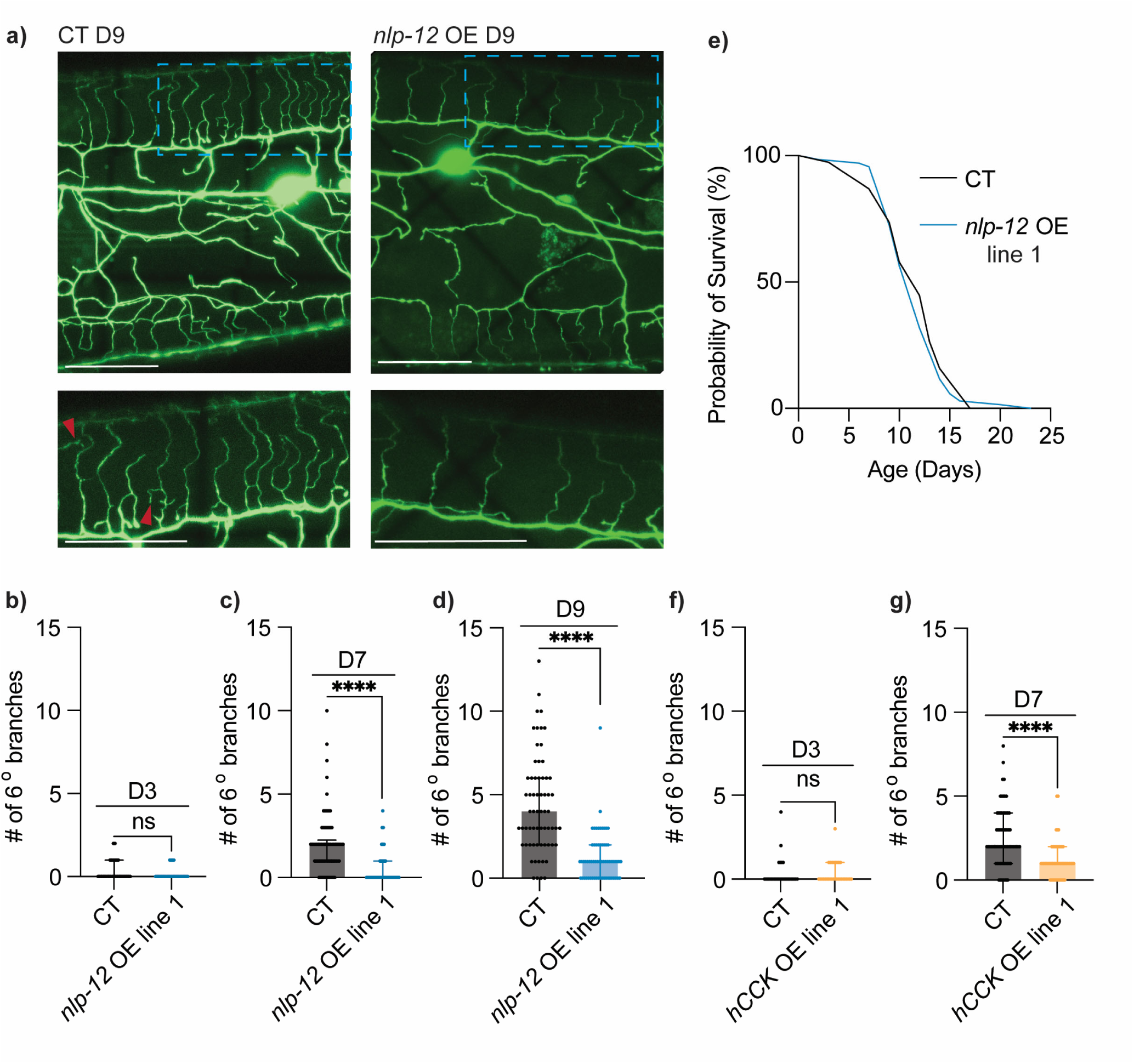
Overexpression of *nlp-12* attenuates aging-associated excessive dendritic branching. **a)** Representative images of PVD neurons in CT and *nlp-12* overexpression (OE) driven by *nlp-12p* (line 1, *lxyEx131*) at D9. Red arrowheads indicate 6° branches. Scale bar = 25 μm. **b-d)** Quantification of 6° dendrites in CT and *nlp-12* OE driven by *nlp-12p* (line 1) at D3 (CT: *n* = 48, *nlp-12* OE: *n* = 54), D7 (CT: *n* = 90, *nlp-12* OE: *n* = 70) and D9 (CT: *n* = 65, *nlp-12* OE: *n* = 63). **e)** Lifespan comparison of CT and *nlp-12* OE driven by *nlp-12p* (line 1) (Starting *n* = 100). No statistically significant difference between CT and OE animals. Lifespan experiment performed without FUDR. **f-g)** Quantification of 6° dendrites in CT and human CCK (*hCCK*) OE driven by *nlp-12p* with *nlp-12* signal peptide (line 1) at D3 (CT: *n* = 30, *hCCK* OE: *n* = 30) and D7 (CT: *n* = 69, *hCCK* OE: *n* = 60). Data from additional replicates and independent transgenic lines are presented in Figure S3. The Mann–Whitney test was used for two group comparisons and survival data was analyzed using the Kaplan–Meier analysis with log-rank (Mantel–Cox) tests. ns—not significant, **** *p* < 0.0001.

Having found that human CCK could substitute for NLP-12 in *nlp-12* mutants, we next asked whether human CCK could confer similar protection when elevated in otherwise wild-type animals. We found that low-dose human CCK expression likewise did not alter PVD 6° branching at D3 but significantly reduced excessive branching at D7 (Fig. 3f-g; Fig. S3c).

Because loss of *nlp-12* affects both PVD morphology and proprioception, we next asked whether the structural protection conferred by *nlp-12* overexpression was accompanied by functional preservation. At D7, *nlp-12* overexpression did not significantly alter mean wavelength, mean amplitude, or the within-individual variability of either measure compared with age-matched controls (Fig. S3d-g). To assess broader locomotor activity during aging, we also measured one-hour travel distance at D5 and D7. Travel distance declined from D5 to D7 in both control and *nlp-12* overexpression animals, and this decline was not attenuated by *nlp-12* overexpression (Fig. S3h-k). Thus, despite protecting PVD dendritic morphology, *nlp-12* overexpression did not detectably preserve either PVD-associated proprioceptive performance or overall locomotor activity under the conditions examined. However, because both behavioral assays integrate broader sensorimotor and neuromuscular function that also decline with age, they may not isolate functional benefits specifically attributable to preserved PVD morphology.

Collectively, these findings establish that elevating either NLP-12 or human CCK is sufficient to restrain aging-associated PVD dendritic remodeling.

### 2.5 Silencing the NLP-12-producing DVA interneuron triggers early-onset PVD excessive dendritic branching

Since NLP-12 is produced predominantly by the DVA interneuron (Bhattacharya et al. 2014; Chen et al. 2024; Li et al. 2006), we next asked whether normal DVA activity is required to restrain aging-associated PVD excessive branching. We first confirmed the expression pattern of *nlp-12* using an endogenous *nlp-12* locus tagged with T2A::3xNLS::GFP, which showed signal restricted to the DVA soma (Fig. 4a).

**Figure 4:**
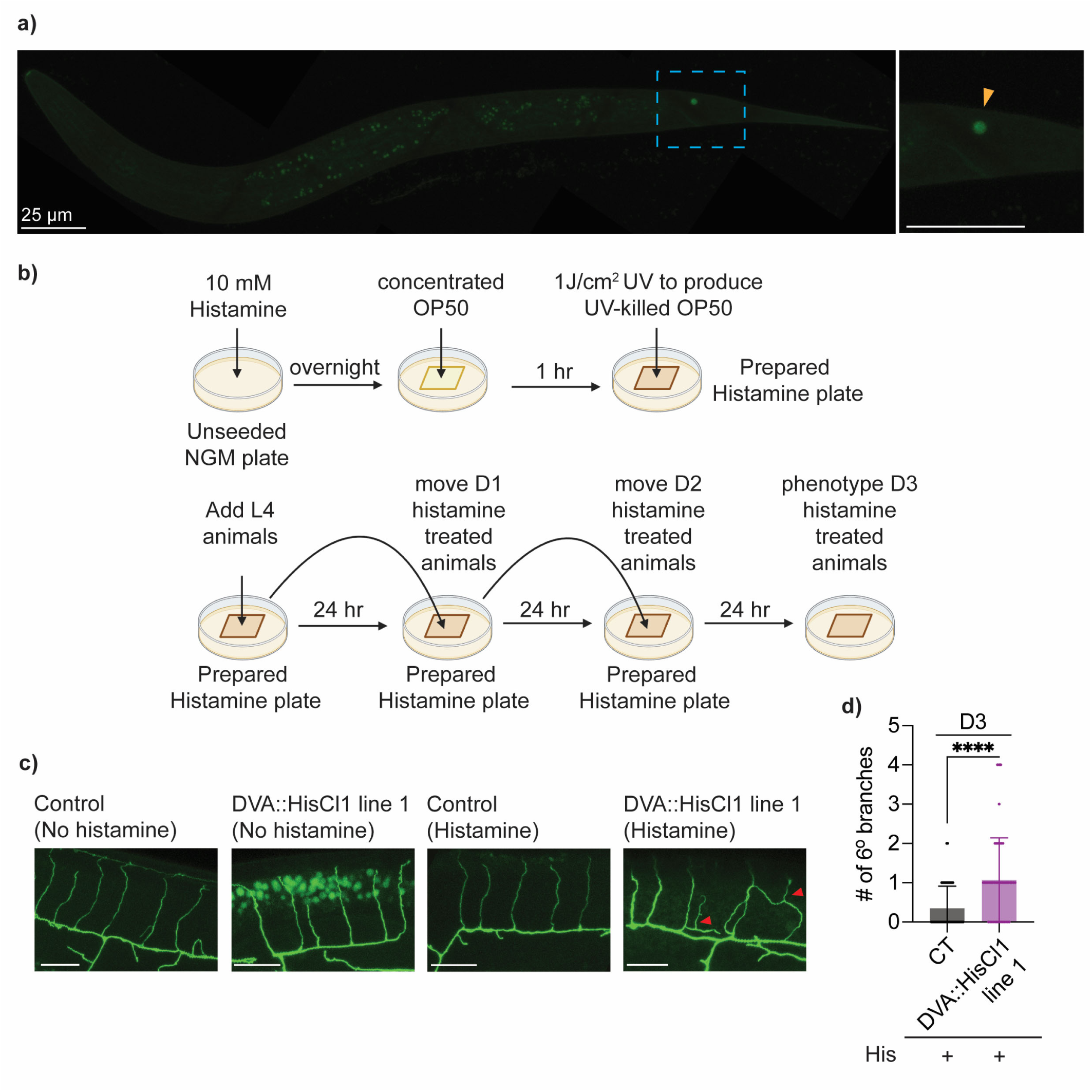
Silencing the NLP-12-producing DVA interneuron triggers early-onset PVD excessive dendritic branching. **a)** Representative image of an L3 stage hermaphrodite with endogenous *nlp-12* locus tagged with T2A::3xNLS::GFP (*syb3112[nlp-12::T2A::3×NLS::GFP]*) demonstrates nuclear GFP signal in the DVA soma. Reporter expression was examined in L4-stage hermaphrodites as well. The yellow arrowhead indicates the location of the DVA soma. Scale bar = 25 μm. Mild autofluorescence of gut granules, a normal physiological feature in *C. elegans*, occasionally appears as small punctate structures throughout the body. **b)** Schematic illustrating histamine plate preparation and treatment protocol (created with BioRender with modifications; https://BioRender.com/mt0fm0w). **c)** Representative images of PVD dendrites in control strain and DVA::HisCl1 animals (line 1; *nlp-12p* driven expression of *Drosophila* histamine-gated chloride channel) at D3, with or without histamine treatment. Each image is centered on a single dendritic menorah. Red arrowheads indicate 6° branches. Scale bar = 10 μm. **d)** Quantification of 6° branches in CT (*n* = 46) and DVA::HisCl1 (line 1) (*n* = 45) at D3. + indicates the presence of histamine (His) on plates from the L4 to D3 stage. Data from additional independent transgenic lines are presented in Figure S4. Experiments performed without FUDR. The Mann-Whitney test was used for two-group comparisons. **** *p* < 0.0001.

To selectively silence DVA, we generated a transgenic strain expressing the *Drosophila* histamine-gated chloride channel HisCl1 under the *nlp-12* promoter, a standard promoter widely used for driving transgene expression and targeted manipulations in the DVA interneuron (Choi et al. 2015; Wen et al. 2012), and treated these animals with exogenous histamine. This chemogenetic approach enables inducible neuronal silencing, as histamine opens HisCl1 to drive chloride influx and suppress neuronal activity (Pokala et al. 2014). Because *C. elegans* lacks endogenous histamine-gated chloride channels, histamine has minimal effects in animals and cells lacking HisCl1 and can therefore selectively inhibit neurons that transgenically express the channel (El Mouridi et al. 2021; Pokala et al. 2014). We initiated histamine treatment at the L4 stage and continued until D3 (Fig. 4b). Under these conditions, we observed that DVA silencing resulted in early-onset excessive dendritic branching by D3 (Fig. 4c-d; Fig. S4a), suggesting that normal DVA activity is required for preserving PVD dendritic integrity during aging.

We next considered whether the *nlp-12* promoter could drive HisCl1 outside DVA in a way that indirectly affects PVD branching. Given that minimal *nlp-12* expression has been reported in the pharynx in addition to DVA (Chen et al. 2024), we considered the possibility that HisCl1 expression driven by the *nlp-12* promoter could also affect pharyngeal activity and thereby alter food intake, which in turn can influence aging-related phenotypes (Lakowski and Hekimi 1998). While we did not directly measure pumping in histamine-treated *nlp-12p::HisCl1* animals, *nlp-12(ok335)* mutants displayed normal pharyngeal pumping under our assay conditions (Fig. S4b), arguing that *nlp-12* does not play a critical role in food intake behavior. In addition, expression of *nlp-12p::HisCl1* did not affect PVD morphology in the absence of histamine, and histamine exposure alone did not alter PVD dendritic morphology in animals lacking HisCl1 as well (Fig. S4c). These controls reduce the likelihood that the branching phenotype results from the transgene alone, histamine exposure, or feeding behavior changes.

Because DVA silencing phenocopied the early-onset PVD branching observed in *nlp-12* mutants, we asked whether it might also affect NLP-12 extracellular delivery. In animals carrying both *nlp-12p::HisCl1* and *NLP-12::mKate*, coelomocyte-associated mKate signal appeared lower following histamine treatment than following vehicle treatment (Fig. S4d). Although this was a qualitative observation, it is consistent with the possibility that DVA activity preserves PVD dendritic integrity in part by supporting NLP-12 secretion.

### 2.6 The cholecystokinin receptors CKR-1 and CKR-2 function downstream of NLP-12 to regulate PVD dendritic integrity during aging

The next step in understanding the mechanism of NLP-12 regulation of PVD neuron aging is to identify potential downstream factors. Because our prior work showed that the apical epidermal collagen *col-120* modulates aging-associated PVD excessive branching in a similar manner (Krishna et al. 2025), we tested its genetic interaction with *nlp-12*. Notably, the severity of excessive branching in *col-120(sy1526)* animals was comparable to that in *nlp-12(ok335)*, and double mutants also did not show enhanced hyperbranching relative to either single mutant (Fig. S5a). Surprisingly, loss of *col-120* completely eliminated the beneficial effects conferred by *nlp-12* overexpression in aged animals (Fig. S5b). These results suggest that epidermal collagen state can gate the phenotypic impact of increased NLP-12 signaling, but do not identify the signaling components that may directly receive and transmit the NLP-12 signal.

We therefore tested the hypothesis that this neuropeptide signal is transmitted through downstream receptors, as neuropeptide ligands typically exert their effects via specific GPCRs. CKR-1 and CKR-2 are both highly conserved homologs of mammalian cholecystokinin receptors and function as receptors for NLP-12 in regulating state-dependent motor patterning and foraging behaviors (Bhattacharya et al. 2014; Chen et al. 2024; Hu et al. 2011; Hu et al. 2015; Peeters et al. 2012; Ramachandran et al. 2021). NPR-2 and NPR-5 are two other GPCRs that share sequence homology with CKR-1 and CKR-2 and have been proposed as potential targets for NLP-12 signaling (Bhattacharya et al. 2014). Upon examining mutants for these receptor candidates, we found that both *ckr-1(yum1505)* and *ckr-2(yum1506)* animals exhibit early-onset excessive PVD dendritic branching at D3, to a similar extent as that observed in *nlp-12(ok335)*, while PVD dendritic morphology was unaffected in *npr-2(ok419)* or *npr-*5*(ok1583)* animals (Fig. 5a-b; Fig. S5c). These results argue that the PVD excessive dendritic branching phenotype is not broadly sensitive to perturbing GPCR signaling, and instead is selectively associated with loss of CKR-1 and CKR-2.

**Figure 5:**
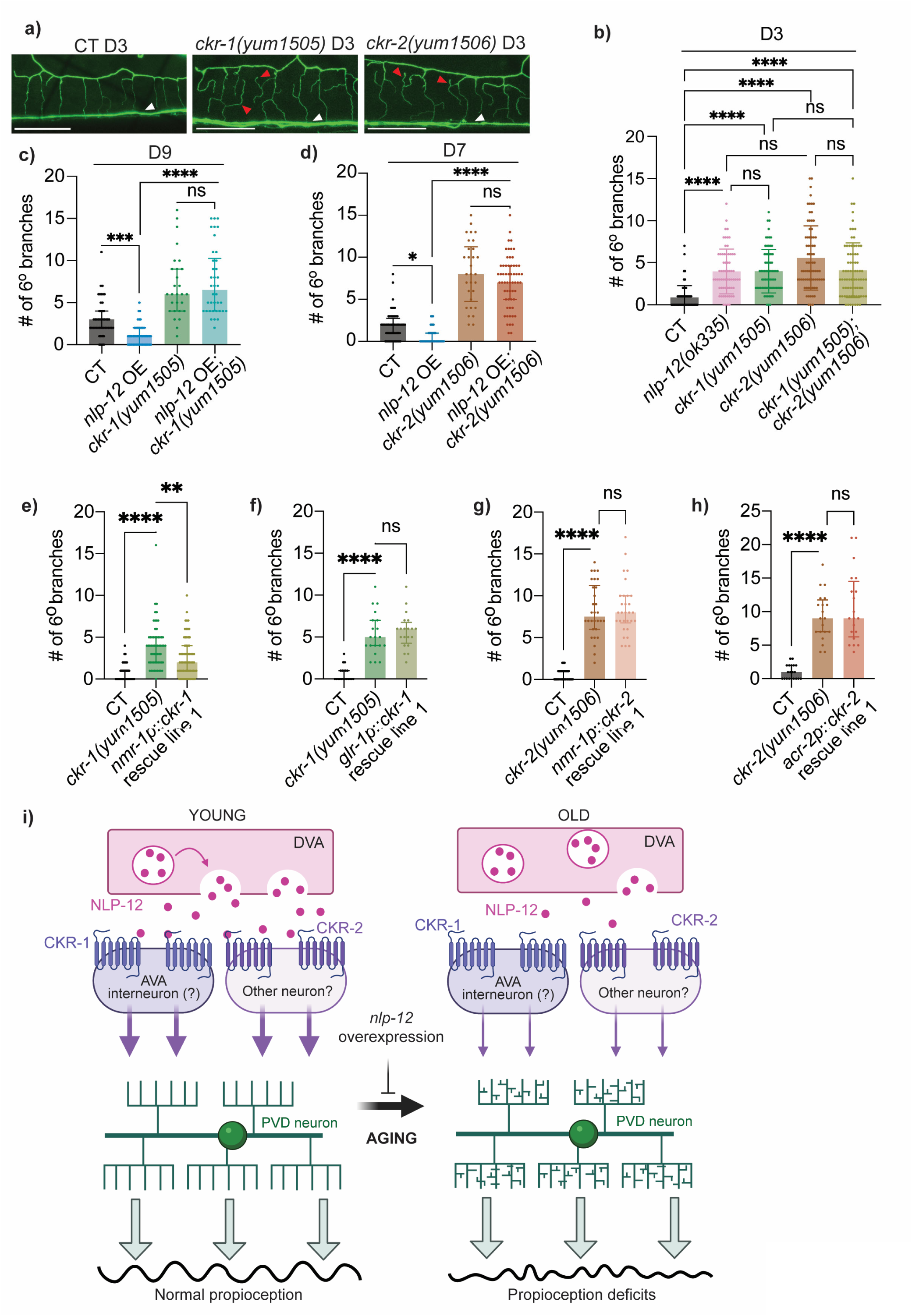
CKR-1 and CKR-2 are required for NLP-12-mediated protection of PVD dendritic health during aging. **a)** Representative images of PVD dendrites in CT, *ckr-1(yum1505)* mutant and *ckr-2(yum1506)* mutant at D3. Each image is centered on a single dendritic menorah. Red arrowheads point to 6° dendritic branches and white arrowheads point to axons. Scale bar = 25 μm. **b)** Quantification of 6° branches in CT (*n* = 77), *nlp-12(ok335)* mutant (*n* = 65), *ckr-1(yum1505)* mutant (*n* = 71), *ckr-2(yum1506)* mutant (*n* = 75) and *ckr-1(yum1505);ckr-2(yum1506)* double mutant (*n* = 77) at D3. **c)** Quantification of 6° branches in CT (*n* = 49), *nlp-12* OE driven by *nlp-12p* (*n* = 50, line 1, *lxyEx131*), *ckr-1(yum1505)* mutant (*n* = 28), and *nlp-12* OE (*lxyEx131*) in the *ckr-1(yum1505)* mutant background (*n* = 38) at D9. **d)** Quantification of 6° branches in CT (*n* = 60), *nlp-12* OE driven by *nlp-12p* (*n* = 40, line 1, *lxyEx131*), *ckr-2(yum1506)* mutant (*n* = 30), and *nlp-12* OE (*lxyEx131*) in the *ckr-1(yum1506)* mutant background (*n* = 60) at D7. **e)** Quantification of 6° branches in CT (*n* = 68), *ckr-1(yum1505)* (*n* = 61), and *ckr-1* rescue driven by *nmr-1p* (line 1) in *ckr-1(yum1505)* (*n* = 66) at D3. **f)** Quantification of 6° branches in CT (*n* = 20), *ckr-1(yum1505)* (*n* = 20), and *ckr-1* rescue driven by *glr-1p* (line 1) in *ckr-1(yum1505)* (*n* = 20) at D3. **g)** Quantification of 6° branches in CT (*n* = 30), *ckr-2(yum1506)* (*n* = 30), and *ckr-2* rescue driven by *nmr-1p* (line 1) in *ckr-2(yum1506)* (*n* = 30) at D3. **h)** Quantification of 6° branches in CT (*n* = 20), *ckr-2(yum1506)* (*n* = 20), and *ckr-2* rescue driven by *acr-2p* (line 1) in *ckr-2(yum1506)* (*n* = 20) at D3. **i)** Working model of NLP-12-dependent regulation of PVD dendritic aging (created with BioRender with modifications; https://BioRender.com/dujlexw). DVA-derived NLP-12 signals through CKR-1 and CKR-2, which may act in distinct downstream neuronal populations, to maintain PVD dendritic integrity. AVA is shown as a candidate site of CKR-1 action based on partial rescue by *nmr-1p::ckr-1*, whereas the site of CKR-2 action remains to be determined. Reduced extracellular NLP-12 availability during aging is proposed to weaken this protective signaling, contributing to PVD hyperbranching and proprioceptive decline, whereas *nlp-12* overexpression delays PVD structural aging. Data from additional independent transgenic lines are presented in Figure S5. The Kruskal–Wallis test with Dunn’s correction was used for multiple-group comparisons. ns—not significant, * *p* < 0.05, ** *p* < 0.01, *** *p* < 0.001, **** *p* < 0.0001.

Notably, *ckr-1;ckr-2* double mutants demonstrated no enhanced branching compared to either single mutant (Fig. 5b), consistent with CKR-1 and CKR-2 functioning in a shared genetic pathway to maintain PVD dendritic integrity. We next tested whether both receptors are required for the protective effect of increased NLP-12 signaling. In older animals, *nlp-12* overexpression attenuates excessive PVD branching in an otherwise wild-type background, but this effect was abolished by either the *ckr-1(yum1505)* or *ckr-2(yum1506)* mutation (Fig. 5c-d). Thus, both CKR-1 and CKR-2 are required downstream of NLP-12 to protect PVD dendritic structure during aging.

Previous reporter studies detected *ckr-1* and *ckr-2* expression in multiple neuronal populations, including interneurons and motor neurons, but did not report detectable expression in PVD (Chen et al. 2024; Hu et al. 2011; Ramachandran et al. 2021) (Fig. S5d). This raised the possibility that these receptors regulate PVD dendritic integrity non-cell-autonomously. We therefore reasoned that receptor-expressing neurons with direct connections to PVD could be candidate sites of action. AVA and PVC are major postsynaptic targets of PVD (Cook et al. 2019; Husson et al. 2012; White et al. 1986), and neuronal single-cell transcriptomic data detect both *ckr-1* and *ckr-2* transcripts in AVA (Taylor et al. 2021) (Fig. S5d).

To begin defining the neuronal sites of receptor action, we performed cell-specific rescue experiments. For CKR-1, we expressed wild-type *ckr-1* under the *nmr-1* promoter, which is active in a restricted set of interneurons that includes AVA and PVC (Brockie et al. 2001; Taylor et al. 2021) (Fig. S5d). Expression of *nmr-1p::ckr-1* partially but significantly reduced the early-onset excessive PVD branching of *ckr-1* mutants (Fig. 5e; Fig. S5e). For comparison, we also expressed *ckr-1* under *glr-1p*, which drives expression in a broader group of command and head neurons, including SMD, a previously established site of CKR-1 function in a stimulus-evoked escape steering behavior (Chen et al. 2022). However, *glr-1p::ckr-1* did not rescue the branching phenotype (Fig. 5f; Fig. S5f).

We next performed analogous rescue experiments for CKR-2. We expressed *ckr-2* under *nmr-1p* to test whether it could function within the same interneuron population that supported partial CKR-1 rescue. We also expressed *ckr-2* under *acr-2p*, which is active in cholinergic ventral cord motor neurons, including VA, VB, DA, and DB neurons, where CKR-2 has previously been shown to mediate NLP-12-dependent potentiation of acetylcholine release at neuromuscular junctions (Hu et al. 2011; Hu et al. 2015). Neither *nmr-1p::ckr-2* nor *acr-2p::ckr-2* was able to rescue the early-onset PVD branching phenotype of *ckr-2* mutants (Fig. 5g-h; Fig. S5g-h).

Collectively, these results establish CKR-1 and CKR-2 as required downstream components of NLP-12-mediated protection of PVD dendritic integrity during aging. Although the cellular site of CKR-2 action remains unresolved, our data support a non-cell-autonomous model of CKR-1 function involving one or more *nmr-1p*-expressing interneurons, with AVA as one candidate (Fig. 5i).

## 3. DISCUSSION

Adult neurons must balance plasticity with stability throughout life, yet the signals that actively restrain aging-associated structural and functional changes *in vivo* remain poorly defined. In this study, using the *C. elegans* PVD sensory neuron as a quantitative readout, we identify conserved CCK-like neuropeptide signaling as an adult maintenance input that buffers an aging-associated dendritic remodeling program. Loss of *nlp-12* accelerates excessive dendritic branching and proprioceptive decline that are typically seen in aged wild-type animals, while restoring the peptide signaling, including by human CCK, suppresses premature changes in dendritic structure. Our receptor genetics indicate that the GPCRs CKR-1 and CKR-2 are required for NLP-12-dependent neuroprotection, highlighting that age-associated neuronal decline may be tuned through a receptor-defined neuromodulatory pathway. Collectively, these findings establish a tractable *in vivo* framework for dissecting how neuromodulatory signals are deployed across adulthood to sustain neuronal healthspan, without necessarily altering organismal lifespan.

Prior work has defined NLP-12 as a DVA-derived CCK-like neuropeptide that signals through the GPCRs CKR-1 and CKR-2 to shape locomotor and foraging-related behaviors (Bhattacharya et al. 2014; Hums et al. 2016; Janssen et al. 2008; Ramachandran et al. 2021). Importantly, this ligand-receptor axis has been anchored to distinct circuit and effector sites. Our work expands this ligand-receptor axis beyond behavioral modulation to an aging-relevant role in adult neuronal maintenance. However, this raises the question of which step becomes limiting with age.

Mechanistically, our results highlight peptide availability as an age-sensitive control point for NLP-12 signaling. While *nlp-12* expression does not show a strong age-linked change, aging is accompanied by reduced coelomocyte-associated NLP-12 signal and increased retention in DVA soma, consistent with diminished effective secretion over time. Since coelomocyte uptake is an indirect proxy for extracellular peptide delivery rather than a direct measurement of release from DVA, it may also be influenced by coelomocyte activity and whole-animal physiology. Nevertheless, the failure of a signal-sequence-mutant *nlp-12* construct to rescue the excessive branching phenotype in *nlp-12* mutants indicates that correct routing through the secretory pathway, and thus extracellular signaling, is required for protection, although signal-sequence disruption may also affect processing or peptide stability. The DVA-silencing findings further raise the possibility that activity-dependent peptide delivery links the functional state of a peptide-producing neuron to the structural maintenance of a distant sensory neuron. Together, these observations suggest that age-related changes in peptide processing, transport, or activity-coupled release could reduce neuropeptide output even when expression persists (Laurent et al. 2018; Sasidharan et al. 2012; Topalidou et al. 2016).

If neuropeptide availability becomes limiting with age, the downstream receptors become the key gate for whether the remaining signal can still be protective. Our findings indicate that both CKR-1 and CKR-2 are required for NLP-12-mediated protection of PVD dendritic integrity. The non-additive phenotype of the *ckr-1;ckr-2* double mutant suggests that the receptors contribute to a shared protective process. Yet, a shared protective process does not require the two receptors to act in the same cells, as neuropeptides can signal extrasynaptically to distributed targets. CKR-1 and CKR-2 have broad, partly overlapping neuronal distributions and have previously been implicated in NLP-12-dependent modulation of distinct components of the motor circuit (Hu et al. 2011; Hu et al. 2015; Ramachandran et al. 2021; Taylor et al. 2021). One possibility is that CKR-1 and CKR-2 act in separate neuronal populations whose outputs converge to establish a circuit, signaling, or mechanical state that supports PVD dendritic maintenance. Alternatively, the receptors may function sequentially at different nodes of the same sensorimotor feedback pathway, such that loss of either receptor disrupts the protective response.

Another important circuit-level question is how DVA-derived NLP-12 influences PVD branching dynamics when neither receptor has been detected in PVD in available expression maps. Although connectome data identified sparse synaptic connectivity between DVA and PVD (Cook et al. 2019), these contacts do not provide an obvious route for the CKR-1- and CKR-2-dependent effects observed here. Instead, available expression and functional mapping place the receptors in multiple interneuron and motor-neuron populations (Hu et al. 2011; Ramachandran et al. 2021; Taylor et al. 2021). Our cell-specific rescue results implicate one or more *nmr-1p*-expressing interneurons in CKR-1 function. Among these neurons, AVA is an intriguing candidate because single-cell transcriptomic data detect *ckr-1* in AVA (Taylor et al. 2021), and AVA receives synaptic input from PVD (Cook et al. 2019). However, because *nmr-1p* is active in multiple interneurons, the precise cellular site of CKR-1 action remains to be established, and the cellular site of CKR-2 action also remains unresolved.

Together, these findings favor a model in which NLP-12 acts through receptor-expressing neurons outside PVD, whose outputs indirectly influence PVD dendritic maintenance.

One plausible way in which this distributed circuit could influence PVD is by altering the pattern of mechanical input that PVD receives during locomotion. Body bending activates DVA, leading to NLP-12 release and modulation of motor output through CKR-1- and CKR-2-dependent pathways that regulate distinct components of head and body motor circuits (Hu et al. 2011; Ramachandran et al. 2021). An age-related decline in NLP-12 availability could therefore alter the timing or coordination of muscle contraction and body bending, changing the repeated deformation and proprioceptive stimulation experienced by PVD dendrites. Because PVD terminal branches run along the body wall at the interface between the epidermis and body-wall muscle and sense body movement (Liang et al. 2015; Tao et al. 2019), persistent changes in this input could plausibly shift the balance between extension and retraction of their dynamic, actin-rich higher-order branches, leading over time to net branch accumulation (Krishna et al. 2025). A key prediction is that restoring appropriate motor-circuit output downstream of NLP-12 should suppress excessive PVD branching in *nlp-12* mutants. The same framework also provides one interpretation of the genetic interaction between *nlp-12* and *col-120*. Changes in the apical epidermal collagen COL-120 may influence the mechanical properties of the epidermal and cuticular scaffold and thereby determine how locomotion-associated deformation is transmitted to PVD (Krishna et al. 2025). Under this model, NLP-12/CKR signaling shapes motor-generated mechanical input, whereas epidermal ECM state influences how that input reaches PVD, allowing circuit and tissue states to jointly regulate dendritic stability during aging.

Our data showing protection of PVD dendritic structure and partial functional rescue by human CCK indicate that key features required for protective CCK signaling are conserved across substantial evolutionary distance. This suggests that the protective effect depends on conserved properties of the ligand-receptor logic rather than *C. elegans*-specific properties of NLP-12. More broadly, studies with human postmortem brains have reported region-dependent alterations in CCK and receptors in Alzheimer’s disease (AD), with some variability across cohorts and assays (Hays and Paul 1982; Lofberg et al. 1996; Mazurek and Beal 1991; Rossor et al. 1981). More recent human biomarker work with the Alzheimer’s Disease Neuroimaging Initiative (ADNI) also reports that cerebrospinal fluid CCK levels are associated with cognitive performance and neuroimaging changes (Plagman et al. 2019). Additionally, studies with AD mouse models further suggest that CCK-linked circuitry and CCKBR, a principal receptor for CCK, can influence disease-relevant synaptic and cognitive phenotypes (Shi et al. 2020; Zhang et al. 2024). These observations link CCK signaling to mammalian neurodegeneration in disease-relevant contexts, but they do not yet establish whether reduced CCK tone is sufficient to drive age-associated neuronal decline. In that light, our results support targeted tests in vertebrate systems of whether CCK signaling contributes to long-timescale neuronal maintenance during aging, and whether altered peptide availability or release, including secretion-dependent changes, contributes to neuronal vulnerability.

In summary, this work identifies conserved CCK-like neuropeptide signaling as a non-cell-autonomous maintenance input that buffers aging-associated neuronal decline at both structural and functional levels in *C. elegans*. It further highlights secretion-dependent neuropeptide availability as a potential age-sensitive bottleneck, suggesting a route by which aging can weaken neuromodulatory maintenance signals even when gene expression is largely preserved. More broadly, together with prior demonstrations that non-neural tissue cues reshape PVD aging (E et al. 2018; Krishna et al. 2025), these findings support a model in which extrinsic signals from multiple origins, including barrier tissues and other neurons, tune selective aspects of neuronal aging.

## 4. MATERIALS AND METHODS

### 4.1 *Caenorhabditis elegans* Strains and Maintenance

*C. elegans* strains were maintained at 20°C, or as otherwise indicated, on Nematode Growth Medium (NGM) plates seeded with *E. coli* OP50. Only hermaphrodites were used for data-generating experiments. For age-synchronized progeny, plates with gravid hermaphrodites were bleached to eliminate microbes, and animals were transferred to NGM plates containing 100 μM 5′fluorodeoxyuridine (FUDR) (VWR, 76345–984) at the mid- or late-L4 stage to prevent egg hatching. Day 1 of adulthood (D1) was defined as 24 hours post-L4. Certain experiments were performed without FUDR, moving animals onto fresh plates daily until D5, and every 2 days thereafter. FUDR-free experiments are noted in figure legends. See Table S1 for strain information.

### 4.2 DNA Constructs and Generation of Transgenes

DNA expression constructs were generated using Gateway cloning technology (Thermo Fisher) (Table S2). Primers were acquired from Millipore and Thermo Fisher (Table S3). Transgenic animals were generated by microinjection, with plasmid DNAs used at 1–100 ng/μL, and co-injection marker *ttx-3p::GFP* or *ttx-3p::RFP* at 50 ng/μL (Tables S1 and S2). At least two independent transgenic lines were analyzed per construct. All *nlp-12*-based transgenes were driven by the *nlp-12* promoter, comprising the 320-bp region immediately upstream of the *nlp-12* translational start codon. The *nlp-12* rescue/overexpression construct encoded untagged *C. elegans* NLP-12 (WormBase WBGene00003750; RefSeq protein NP_490908.1) from genomic sequence followed by the native *nlp-12* 3′UTR. The NLP-12 secretion reporter encoded a direct in-frame C-terminal translational fusion of NLP-12 to mKate, driven by *nlp-12p* and followed by the native *nlp-12* 3′UTR; no linker sequence was inserted. The human CCK rescue construct was generated from a human CCK ORF clone (Applied Biological Materials, PC115339; RefSeq mRNA NM_000729.6; RefSeq protein NP_000720.1). The endogenous N-terminal signal peptide of human CCK was removed and replaced with the *C. elegans nlp-12* genomic region encoding the NLP-12 signal peptide (residues 1-20; NP_490908.1), including its native intronic sequence. This NLP-12 signal-peptide-encoding region was joined directly and in frame, without an intervening linker, to the human CCK coding sequence lacking its native signal peptide. The resulting construct was driven by *nlp-12p* and followed by the native *nlp-12* 3′UTR. Cell-specific receptor-rescue constructs encoded full-length, untagged *ckr-1* or *ckr-2* cDNA. The *ckr-1* cDNA (WormBase WBGene00020712) was expressed under either *nmr-1p* or *glr-1p*, whereas the *ckr-2* cDNA (WormBase WBGene00021439) was expressed under either *nmr-1p* or *acr-2p*. The *nmr-1*, *glr-1*, and *acr-2* promoter fragments comprised 1,104 bp, 755 bp, and 1,743 bp immediately upstream of their respective translational start codons. For the DVA-silencing constructs, HisCl1 and mScarlet were separated by an *SL2* trans-splicing element, allowing expression of the two proteins as separate gene products from the same transgene, with mScarlet serving as a fluorescent marker of transgene expression while HisCl1 functions as the silencing channel. pHW1385 (*myo-2p::HisCl::SL2::mScarlet::tbb-2 3’UTR*) was a gift from Dr. Han Wang (Addgene plasmid # 229879) (Khan et al. 2025).

### 4.3 Fluorescence Microscopy

Neurons were visualized using fluorescent reporters and phenotyped on a Zeiss Axio Imager M2 compound microscope using 5 mM of levamisole (Millipore Sigma, 31742) for immobilization and mounted on 4% agarose pads in M9 solution. Confocal images were taken on a Leica SP8 confocal microscope with Z-stack (0.5-1 μm/slice) at 63x magnification. Imaging parameters were kept constant across genotypes within each experiment.

### 4.4 Quantification of PVD aging phenotypes

*F49H12.4::GFP(wdIs51)* (Smith et al. 2010) was used to visualize PVD neurons. Excessive dendritic branching was quantified on a Zeiss Axio Imager M2 compound microscope by counting 6° branches, identified within “menorah” structures. 6° branches for each animal were counted for either PVDL or PVDR for either the dorsal or ventral side, depending on which was clearly visible during the imaging session. For *mec-10* and *del-1* mutants, *ser-2(prom3)::myr::GFP + odr-1p::RFP(wyIs592)III* was utilized to visualize PVD neurons due to challenges in crossing *mec-10(tm1552)X* and *del-1(ok150)X* with (*wdIs51)X*. *wyIs592* animals had a larger body size than *wdIs51*, likely due to genetic background differences, which may account for the branching number variation at D3 between the two reporter strains.

For each experimental group, a minimum of three replicates of 20 animals each, or as indicated otherwise, were quantified alongside a control group, with experimenters blinded to sample identity to minimize bias.

### 4.5 NLP-12::mKate Imaging

Levels of secreted and retained NLP-12 were assessed via imaging on a Leica SP8 confocal microscope using 2% agarose pads with 5 mM of levamisole for immobilization. Experimental animals [*unc-122p::GFP(oxIs253); NLP-12::mKate; ttx3p::GFP* (*lxyEx120)*] were imaged longitudinally. Each individual animal was imaged at D3, recovered after imaging, and the same individual animal was re-imaged at D7. Thus, the D3 and D7 fluorescence measurements represent paired measurements from the same animal. Images were Z-stack projections captured with a 63x objective in both red and green channels. A total of 3 to 4 sections were imaged for each animal to capture the three coelomocyte pairs and the DVA neuron soma. Fiji was utilized to obtain the mean gray values (MGV) of mKate fluorescence within individual coelomocytes by selecting the area of the coelomocyte as indicated by the coelomocyte::GFP label. The MGVs were averaged across all coelomocytes within each animal to obtain a representative coelomocyte MGV for each animal (Fig. 2c) or summed up to obtain the total MGV for each animal (Fig. S2b) to estimate NLP-12 within all coelomocytes. NLP-12::mKate levels in the DVA were measured by obtaining the MGV of mKate fluorescence within the DVA soma region. The localization of this signal to the DVA cell body was independently confirmed by crossing the NLP-12::mKate reporter with ZM11177 *[hpIs860; twk-40p(short)::eGFP]*, which labels DVA with cytosolic GFP, and examining overlap between the GFP-labeled DVA soma and NLP-12::mKate fluorescence (Fig. S2c).

### 4.6 Lifespan Assay

Lifespan assays were conducted at 20 °C in the absence of FUDR. L4 animals were transferred to assay plates and scored daily starting from Day 1 of adulthood. Animals were transferred daily until Day 5 of adulthood and then every other day thereafter to minimize confounding from progeny.

Lifespan was calculated from the time animals were first placed on the assay plates until they were scored as dead. An animal was considered dead when it failed to respond to gentle prodding with a platinum wire and displayed no pharyngeal pumping. Animals that died due to vulval rupture, internal hatching (bagging), or desiccation (crawling off the agar) were censored from the analysis. Each condition included 90–100 animals at the starting points across three biological replicates. The PVD::GFP background strain (*wdIs51*) was used as the control for all experiments.

### 4.7 PVD Function Analysis: Proprioception

Proprioceptive sensing specific to PVD neurons was assessed using a locomotion assay as described previously (Krishna et al. 2025; Tao et al. 2019; Tavernarakis et al. 1997) with some modifications. On the day of the assay, animals were transferred to a fresh NGM plates seeded with OP50, with one animal per plate, which was allowed to roam freely for 2–3 h at 20°C, until sufficient tracks were made in the bacterial lawn. The plate was then imaged using a digital camera attached to a Leica S9i stereomicroscope at 1x magnification. A total of 100 randomly selected wavelengths (distance between two adjacent peaks) and amplitudes (distance between peak and wavelength, perpendicular to wavelength) were measured using Fiji. All measurements were used in cases where animals had fewer than 100 measurements, with a minimum of 40 measurements per animal. Within-individual variability in wavelength and amplitude was calculated for each animal as the coefficient of variation (CV), defined as the standard deviation divided by the mean of the measurements from that individual animal, multiplied by 100. All measurements were normalized to the body lengths of individual animals and units were converted to millimeters utilizing a calibration slide. All animals subjected to the proprioception assay were also examined under the Zeiss to quantify PVD dendritic branching.

### 4.8 Spontaneous Locomotor Activity Assay

Spontaneous locomotor activity was assessed using a track-based locomotion assay modified from previous studies (Narasimhan et al. 2011). Individual animals were placed in the center of fresh NGM plates seeded with OP50, with one animal per plate, and were allowed to move freely for 1 h at 20°C. Movement tracks were imaged using a Leica S9i stereomicroscope at 0.6x magnification. The cumulative distance traveled by each animal was quantified by tracing the complete movement track using the segmented-line tool in Fiji. Measurements were converted to millimeters using a calibration slide imaged under the same microscope settings. The same animals were assayed longitudinally on D5 and again on D7.

### 4.9 Histamine-based DVA silencing

A standard histamine treatment protocol (Pokala et al. 2014) was utilized, with 1 M histamine solutions prepared by adding 1.85 g of histamine dihydrochloride (Sigma-Aldrich, H7250-10G) to 10 mL of ultrapure water and filtered. 150 μL of 1 M His was added to unseeded 60 mm NGM plates (final concentration of 10 mM) and left to rest overnight. Control plates were prepared with an equal volume of ultrapure water. OP50 was cultured for 18-20 hours in 50 mL of B-Broth and centrifuged, the supernatant was then discarded, and 50 µL of concentrated OP50 was pipetted and spread onto the prepared plates. After drying for 1 hour, the bacteria were killed using UV in a Spectroline Microprocessor-Controlled UV Crosslinker at 1J/cm^2^. UV-killed OP50 was used to prevent bacterial proliferation and histamine-dependent changes in lawn density or bacterial physiology during the multi-day exposure period, which may have altered the feeding/nutrient conditions and confound interpretation of histamine-driven HisCl1 silencing. L4 animals were then added to the prepared plates and moved on to fresh plates every 24 hours until phenotyped at D3. Animals expressing the *Drosophila* HisCl1 channel in pharyngeal muscles [*F49H12.4::GFP(wdIs51); myo-2p::HisCl1::SL2::mScarlet::tbb-2 5’UTR(lxyEx134)*] were used as a positive control to confirm the effectiveness of the prepared histamine plates on all days of the protocol, assessed by examining the pharyngeal pumping of treated positive controls.

### 4.10 Pharyngeal Pumping Assay

To assess pharyngeal pumping in *nlp-12* mutants, healthy actively moving animals were chosen and a Leica S9i stereomicroscope was used to capture individual video recordings of pharyngeal pumping of each animal using a 2x objective lens with brightfield illumination. As the animals were unrestrained and freely moving, the plates were placed on a transparent plastic sheet and moved to keep the animal in the lens’ field of view. The captured video was viewed on VLC media player at 0.25x speed and the number of pharyngeal pumps was counted using a handheld counter for a period of 30 seconds. Animals that were harmed while picking (aberrant track waveforms, unmoving animals, injured animals) and animals not on the bacterial lawn were excluded.

### 4.11 RT-qPCR

To quantify *nlp-12* expression levels in aging, N2 wild-type animals at L4 and D7 were collected, using the 10-worm lysis method (Ly et al. 2015) to prepare lysates. Briefly, 10 animals were cleaned in ultrapure H_2_O and placed in 1 μL of lysis buffer (0.25 mM EDTA, 5 mM Tris pH 8, 0.5% Triton X-100, 0.5% Tween-20, 1 mg/mL proteinase K). Samples were lysed in a thermal cycler at 65°C for 15 min, 85°C for 1 min, and then stored at −80°C for at least 12 h. An additional 1.5 μL of lysis buffer was then added, and the cycle was repeated. Samples were treated with dsDNAse (Thermo Fisher; 0.5 μL each of 10X buffer and dsDNAse, 1.5 μL ultrapure H_2_O) and incubated at 37°C for 15 min, 55°C for 5 min, and 65°C for 1 min, followed by reverse transcription. Reverse transcription was performed with iScript Reverse Transcription Supermix for RT-qPCR (BioRad). RT-qPCR was performed using designed primers (Table S3) and iTaq Universal SYBR Green (Bio-Rad). *cdc-42* and *act-2* were used as housekeeping controls. Primer efficiency was verified utilizing standard curve method. The Bio Rad CFX96 Touch Deep Well Real-time PCR Detection System was used to perform the qPCR and relative mRNA levels were calculated using the comparative ΔΔCT method.

### 4.12 Statistical Analysis

All analyses were performed using GraphPad Prism (v11.0.0). Normality was assessed using the Shapiro–Wilk and D’Agostino & Pearson tests. Survival data were evaluated using Kaplan–Meier analysis with log-rank (Mantel–Cox) tests. The Pearson’s correlation test was applied for correlation analyses. For multiple group comparisons, one-way ANOVA with Tukey’s post hoc test was used for normally distributed datasets, and the Kruskal–Wallis tests with Dunn’s correction were used for non-parametric datasets. For two group comparisons, unpaired two-tailed *t*-tests with Welch’s correction for normally distributed data or the Mann–Whitney tests for non-parametric data were used. Within-animal comparison of NLP-12::mKate levels between D3 and D7 and cumulative distance traveled between D5 and D7 were analyzed using the Wilcoxon matched-pairs signed rank tests. Significance was defined as *p* < 0.05. Bar graphs display the mean ± standard error of the mean (SEM) for normally distributed data, and the median with interquartile range for non-normally distributed data, unless otherwise specified. Sample sizes and statistical tests are provided in the corresponding figure legends.

## DATA AVAILABILITY

Strains and plasmids are available upon request. The authors affirm that all data necessary for confirming the conclusions of the article are present within the article, figures, and supplementary materials.

## ACKNOWLEDGMENTS

Some strains were provided by the CGC, which is funded by NIH Office of Research Infrastructure Programs (P40 OD010440). We also thank Dr. Oliver Hobert for kindly sharing the strain PHX3112 for imaging the endogenous expression pattern of *nlp-12*. Plasmid pHW1385 was a gift from Dr. Han Wang.

## AUTHORS’ CONTRIBUTIONS

L.E and M.M.K. conceived the project; M.M.K. and L.E designed the experiments; M.M.K., S.G.W., G.P.P., E.C.M. and T.S. performed the experiments and collected the data; M.M.K., S.G.W., E.C.M., T.S. and L.E analyzed and interpreted the data; M.M.K. and L.E drafted the manuscript; M.M.K., S.G.W., G.P.P. and L.E edited the manuscript.

## Supplemental Materials

### Supplemental Figure

**Figure S1:**
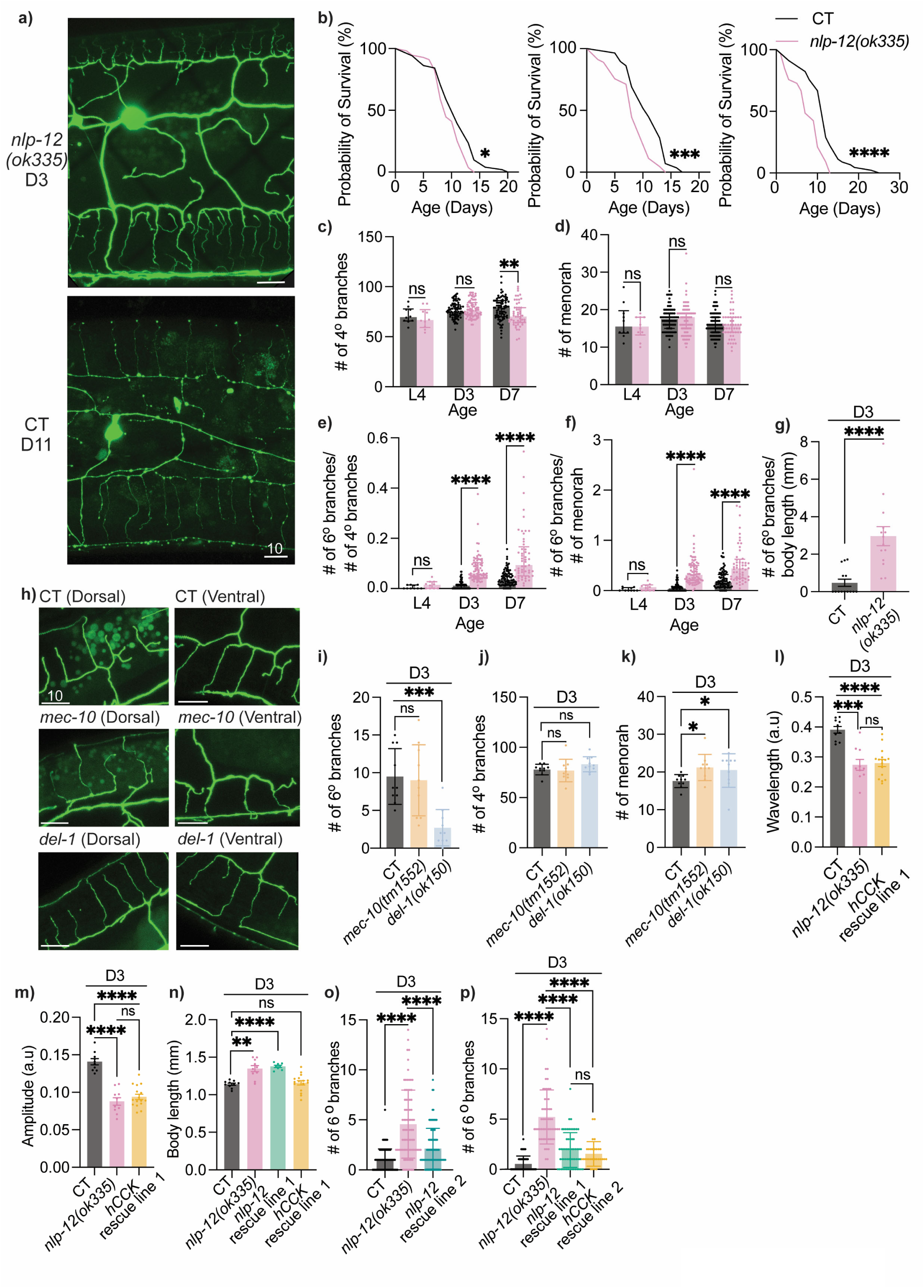
Characterization of NLP-12 regulation of PVD aging. **a)** Representative images showing the absence of early-onset PVD dendritic beading in *nlp-12(ok335)* at D3, compared with CT animals at D11. Scale bar = 10 μm. **b)** Lifespan comparison of CT and *nlp-12* mutants (starting *n* = 100). Performed without FUDR. **c-d)** Quantification of 4° dendrites *(c)* and menorahs *(d)* in CT and *nlp-12* mutants during aging (L4 CT: *n* = 10, L4 *nlp-12* mutant: *n* = 10, D3 CT: *n* = 80, D3 *nlp-12* mutant: *n* = 75, D7 CT: *n* = 82, D7 *nlp-12* mutant: *n* = 53). Dark grey: CT; light pink: *nlp-12(ok335)* mutant. **e-f)** Quantification of 6° dendrites normalized to number of 4° branches *(e)* and menorahs *(f)* in CT and *nlp-12* mutants during aging (L4 CT: *n* = 10, L4 *nlp-12* mutant: *n* = 10, D3 CT: *n* = 80, D3 *nlp-12* mutant: *n* = 75, D7 CT: *n* = 82, D7 *nlp-12* mutant: *n* = 53). Dark grey: CT; light pink: *nlp-12(ok335)* mutant. **g)** Comparison of 6° branches normalized to body length between CT and *nlp-12* mutants at D3 (*n* = 14 each). **h)** Representative images of PVD dendrites in CT, *mec-10(tm1552)*, and *del-1(ok150)* animals at D3. Each image is centered on a single dendritic menorah. Scale bar = 10 μm. **i-k)** Quantification of 6° branches *(i)*, 4° branches *(j)* and menorahs *(k)* in CT, *mec-10(tm1552)* and *del-1(ok150)* animals at D3 (*n* = 10 each). **l-m)** Mean wavelength *(l)* and mean amplitude *(m)* measured from 100 tracks per animal and normalized to body length (CT: *n* = 10, *nlp-12* mutant: *n* = 10, human cholecystokinin/hCCK rescue (driven by *nlp-12p* with *nlp-12* signal peptide, line 1) in *nlp-12(ok335)*: *n* = 16) at D3. **n)** Comparison of body length among CT (*n* = 10), *nlp-12* mutants (*n* = 10), *nlp-12* rescue line 1 (*n* = 8), and human CCK/hCCK rescue line 1 (*n* = 16) at D3. **o)** Quantification of 6° dendrites in CT (*n* = 89), *nlp-12* mutants (*n* = 79), and *nlp-12p::nlp-12* rescue in *nlp-12(ok335)* (line 2) (*n* = 89) at D3. **p)** Quantification of 6° dendrites in CT (*n* = 50), *nlp-12* mutants (*n* = 56), *nlp-12p::nlp-12* rescue in *nlp-12(ok335)* (line 1) (*n* = 58) and hCCK rescue in *nlp-12(ok335)* (driven by *nlp-12p* with *nlp-12* signal peptide, line 2) (*n* = 30) at D3. Survival data were analyzed using the Kaplan–Meier analysis with log-rank (Mantel–Cox) tests. For two group comparisons, unpaired two-tailed *t*-tests with Welch’s correction or the Mann–Whitney tests were used. For multiple-group comparisons, the Kruskal–Wallis tests with Dunn’s correction were used. ns—not significant, * *p* < 0.05, ** *p* <0.01, *** *p* < 0.001, **** *p* < 0.0001.

**Figure S2:**
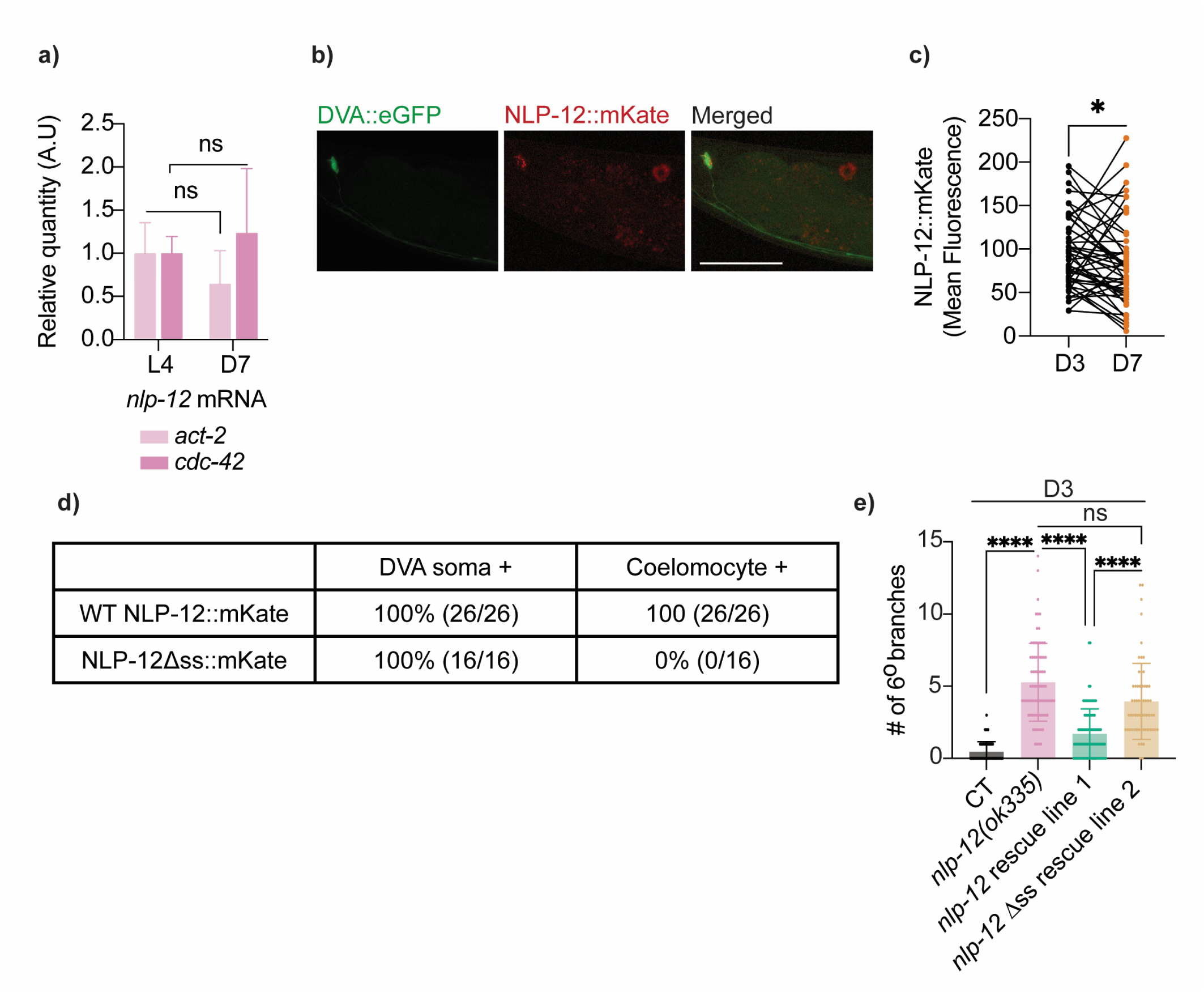
NLP-12 secretion is important for its role in regulating PVD aging. **a)** qPCR analysis of *nlp-12* mRNA expression levels at L4 and D7 in wild-type animals represented as relative quantity, with *act-2* and *cdc-42* as housekeeping genes. **b)** Representative images of an L4 hermaphrodite animal coexpressing NLP-12::mKate and cytosolic GFP in DVA *[twk-40p(short)::eGFP]*. The upper-left NLP-12::mKate signal overlaps with the GFP-defined DVA cell body, whereas the spatially separate red signal in the upper-right is coelomocyte-associated NLP-12::mKate. Scale bar = 50 μm. **c)** Quantification of mean gray value of NLP-12::mKate fluorescence summed across all coelomocytes in the same animals at D3 and D7 (*n =* 48 individual animals measured longitudinally at both ages; each connecting line represents paired measurements from the same individual animal). CT animals *[unc-122p::GFP, nlp-12p::nlp-12::mKate]* were used. *unc-122p::GFP* was used to visualize coelomocytes. **d)** Percentage of L4 animals with detectable NLP-12::mKate fluorescence in the DVA soma and coelomocytes. Wild-type NLP-12::mKate (ELZ287, line 1; *n* = 26) and NLP-12Δss::mKate (ELZ306, line 1; *n* = 16) animals were scored for the presence or absence of detectable mKate fluorescence in each region. The frequency of detectable coelomocyte fluorescence differed significantly between reporters (Fisher’s exact test, *p* < 0.0001). **e)** Quantification of 6° dendrites in CT (*n* = 76), *nlp-12* mutants (*n* = 81), *nlp-12p::nlp-12* rescue in *nlp-12(ok335)* (line 1) (*n* = 86), and *nlp-12 Δss* (mutated signal sequence) rescue (driven by *nlp-12p*, line 2) (*n* = 60) in *nlp-12(ok335)* at D3. The Mann–Whitney test was used for two-group comparisons, the Wilcoxon matched-pairs signed rank test was used for paired comparisons, and the Kruskal–Wallis test with Dunn’s correction was used for multiple-group comparisons. ns—not significant, * *p* < 0.05, **** *p* < 0.0001.

**Figure S3:**
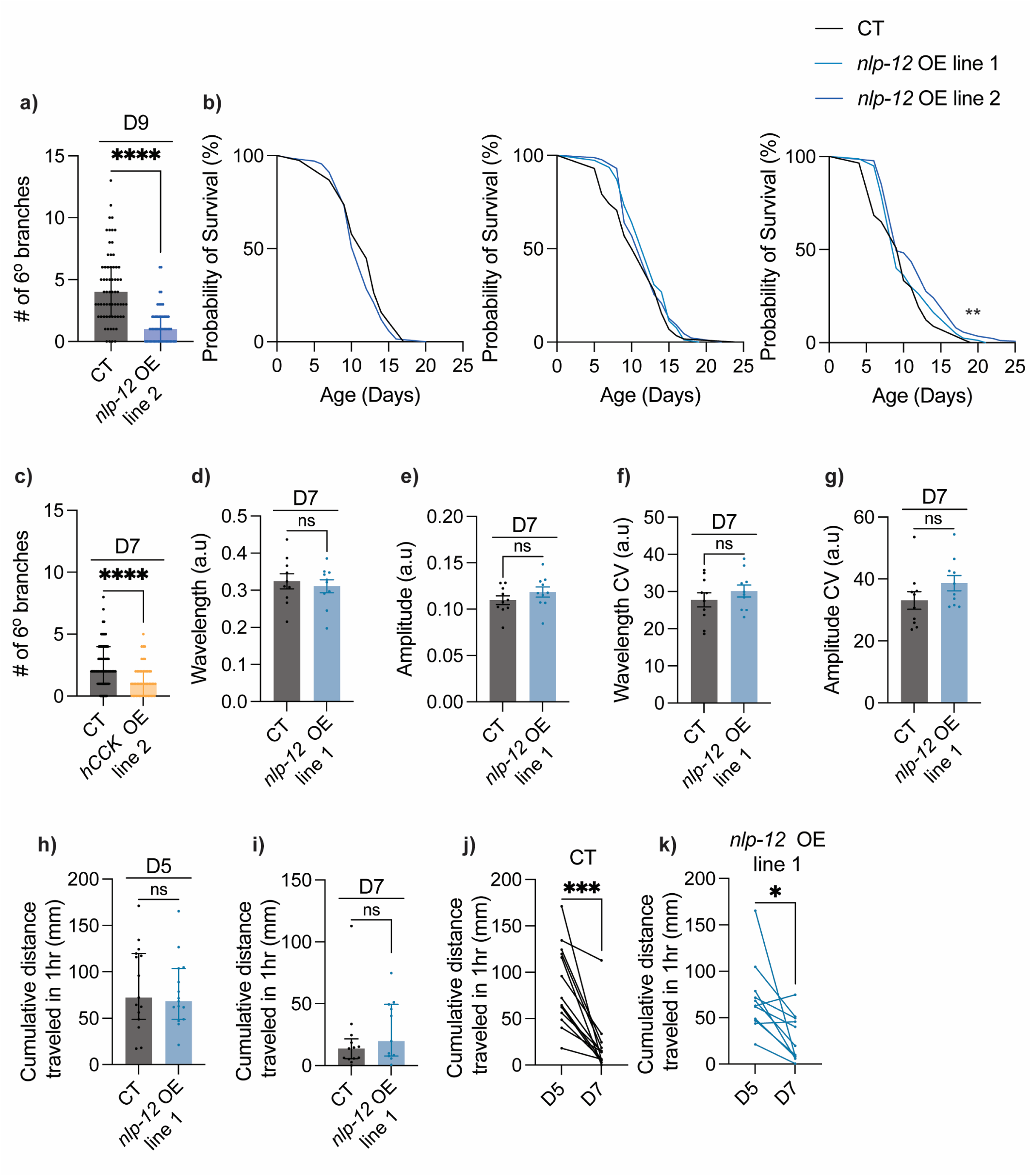
Overexpression of *nlp-12* is neuroprotective. **a)** Quantification of 6° dendrites in CT (*n* = 65) and *nlp-12* overexpression (OE) driven by *nlp-12p* (line 2) (*n* = 63) at D9. **b)** Lifespan comparison of CT and *nlp-12* OE driven by *nlp-12p* (line 1 and line 2) (starting *n* = 100). No statistically significant difference between CT and OE animals unless indicated. Lifespan experiment performed without FUDR. **c)** Quantification of 6° dendrites in CT and human CCK (hCCK) OE driven by *nlp-12p* with *nlp-12* signal peptide (line 2) at D7 (CT: *n* = 69, hCCK OE: *n* = 51). **d-e)** Mean wavelength *(d)* and mean amplitude *(e)* measured from 100 tracks per animal and normalized to body length (CT: *n* = 10, *nlp-12* OE line 1: *n* = 10) at D7. **f-g)** Within-individual variability in wavelength *(f)* and amplitude *(g)* measured via coefficient of variation (CV) from 100 tracks per animal (CT: *n* = 10, *nlp-12* OE line 1: *n* = 10) at D7. **h-k)** Distance traveled during a 1h locomotion assay by control (CT) (*n* = 13) and *nlp-12* OE (line 1) (*n* = 11) animals. (h, i) Between-group comparison at D5 *(h)* and D7 *(i)*. (j, k) Longitudinal comparisons of the same CT *(j)* and *nlp-12* OE *(k)* animals tested at D5 and again at D7. Lines connect measurements from the same individual animal. The proprioception/locomotion experiments were performed without FUDR. Between-group comparisons were analyzed using the Mann–Whitney test, and longitudinal comparisons were analyzed using the Wilcoxon matched-pairs signed-rank test. Survival data was analyzed using the Kaplan–Meier analysis with log-rank (Mantel–Cox) tests. ns—not significant, * *p* < 0.05, *** *p* < 0.001, **** *p* < 0.0001.

**Figure S4:**
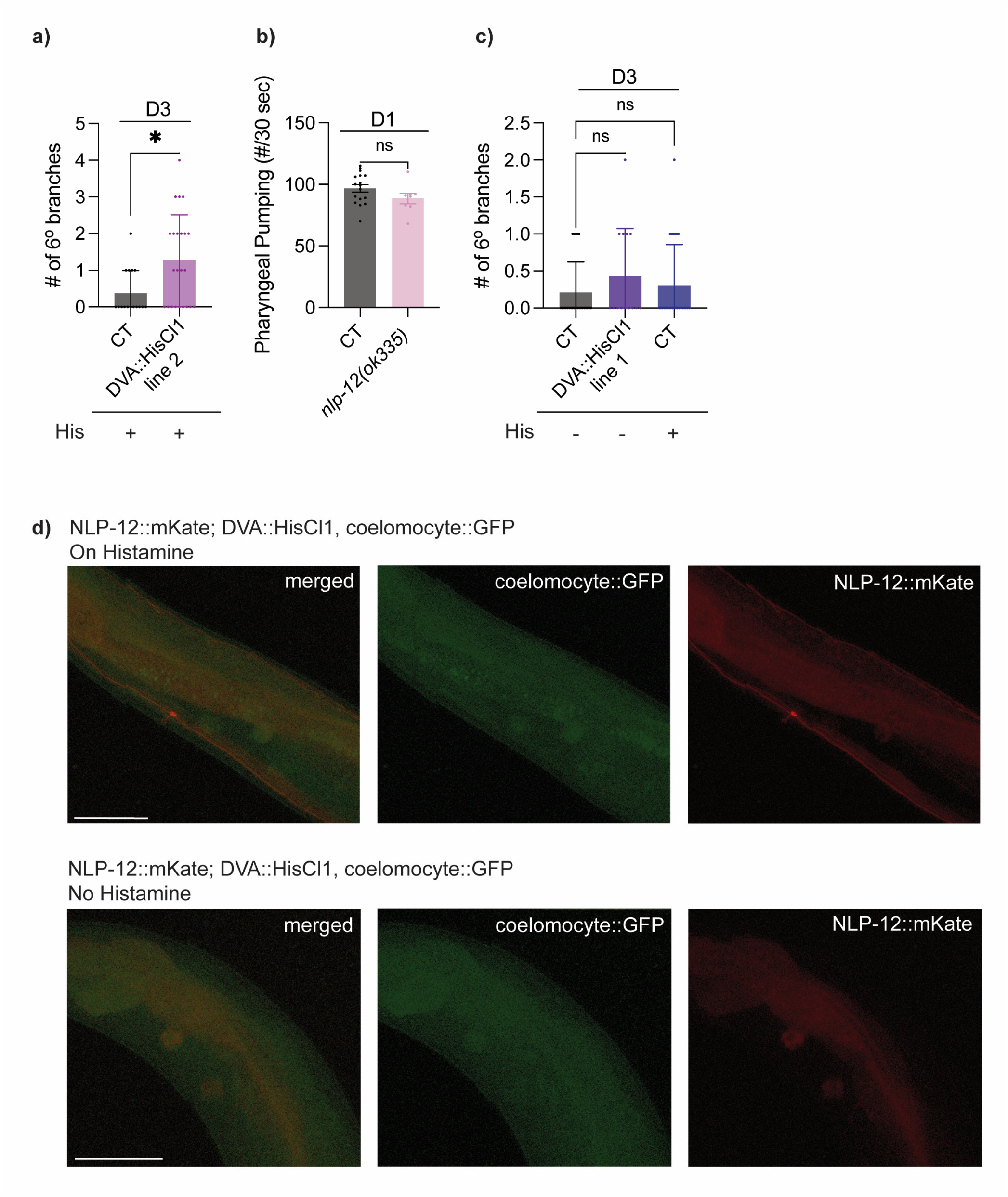
Histamine-based silencing of DVA neuron. **a)** Quantification of 6° branches in CT (*n* = 16) and DVA::HisCl1 (*nlp-12p* driven expression of *Drosophila* HisCl1 channel, line 2, *n* = 23) at D3. **b)** Comparison of pharyngeal pumping rate in CT (*n* = 17) and *nlp-12* mutants (*n* = 8) during a 30 second window at D1. **c)** Quantification of 6° branches in CT (untreated *n* = 24, treated with histamine *n* = 26) and DVA::HisCl1 (line 1, untreated, *n* = 14) at D3. – indicates absence and + indicates presence of histamine (His) on plates from the L4 stage to D3. **d)** Representative images of animals coexpressing the NLP-12::mKate secretion reporter, DVA-silencing transgene *nlp-12p::HisCl1::SL2::mScarlet*, and a coelomocyte GFP marker (ELZ317), with (top) or without (bottom) histamine treatment. The coelomocyte-associated NLP-12::mKate signal appeared lower following histamine-induced DVA silencing. Images are presented qualitatively and were not quantified because the transgenes were carried on independent extrachromosomal arrays. Scale bar = 50 μm. Experiments performed without FUDR. The Mann-Whitney test was used for two-group comparisons and the Kruskal–Wallis test with Dunn’s correction was used for multiple-group comparisons. ns—not significant, * *p* < 0.05.

**Figure S5:**
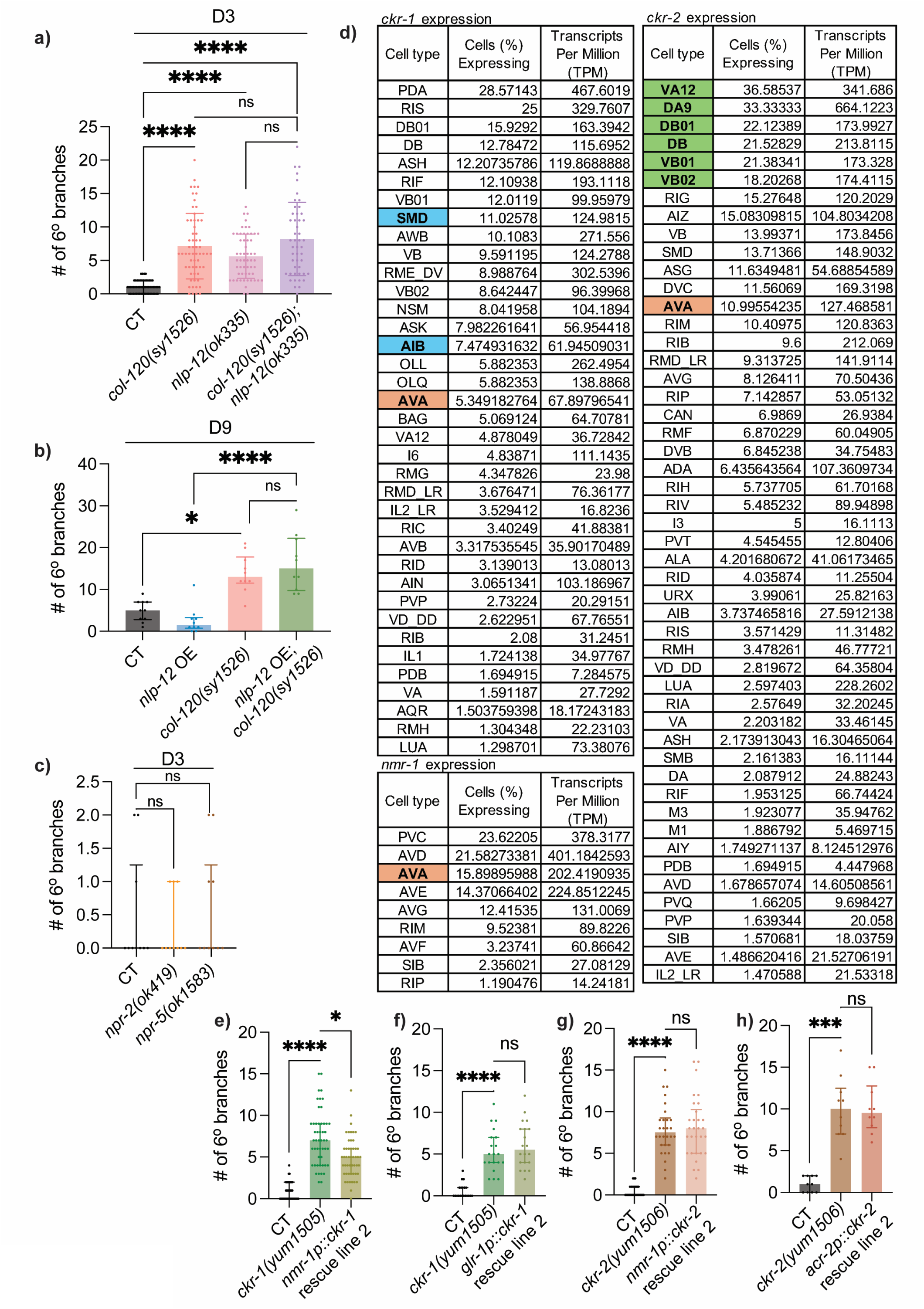
Downstream mechanisms of NLP-12 in regulating PVD aging. **a)** Quantification of 6° branches in CT (*n* = 59), *col-120(sy1526)* (*n* = 60), *nlp-12(ok335)* (*n* = 56), and *col-120(sy1526);nlp-12(ok335)* (*n* = 48) animals at D3. **b)** Quantification of 6° branches in CT, *nlp-12* OE (driven by *nlp-12p*, line 1, *lxyEx131*), and *nlp-12* OE (line 1, *lxyEx131*) in *col-120(sy1526)* background at D9 (*n* = 10 each). **c)** Quantification of 6° branches in CT, *npr-2(ok419)* mutants, and *npr-5(ok1583)* mutants at D3 (*n* = 10 each). **d)** Single-cell transcriptomic expression of *ckr-1*, *ckr-2*, and *nmr-1* across neuronal cell types in wild-type L4 *C. elegans*. Data were retrieved from WormBase and derived from the CeNGEN dataset. **e)** Quantification of 6° branches in CT (*n* = 40), *ckr-1(yum1505)* (*n* = 50), and *ckr-1* rescue driven by *nmr-1p* (line 2) in *ckr-1(yum1505)* (*n* = 50) at D3. **f)** Quantification of 6° branches in CT (*n* = 20), *ckr-1(yum1505)* (*n* = 20), and *ckr-1* rescue driven by *glr-1p* (line 2) in *ckr-1(yum1505)* (*n* = 20) at D3. **g)** Quantification of 6° branches in CT (*n* = 30), *ckr-2(yum1506)* (*n* = 30), and *ckr-2* rescue driven by *nmr-1p* (line 2) in *ckr-2(yum1506)* (*n* = 30) at D3. **h)** Quantification of 6° branches in CT (*n* = 10), *ckr-2(yum1506)* (*n* = 10), and *ckr-2* rescue driven by *acr-2p* (line 2) in *ckr-2(yum1506)* (*n* = 10) at D3. The Mann-Whitney test was used for two group comparisons, and the Kruskal–Wallis test with Dunn’s correction was used for multiple group comparisons. ns—not significant, * *p* < 0.05, *** *p* < 0.001, **** *p* < 0.0001.

## Supplemental Materials

### Supplemental Tables

**Table S1:**
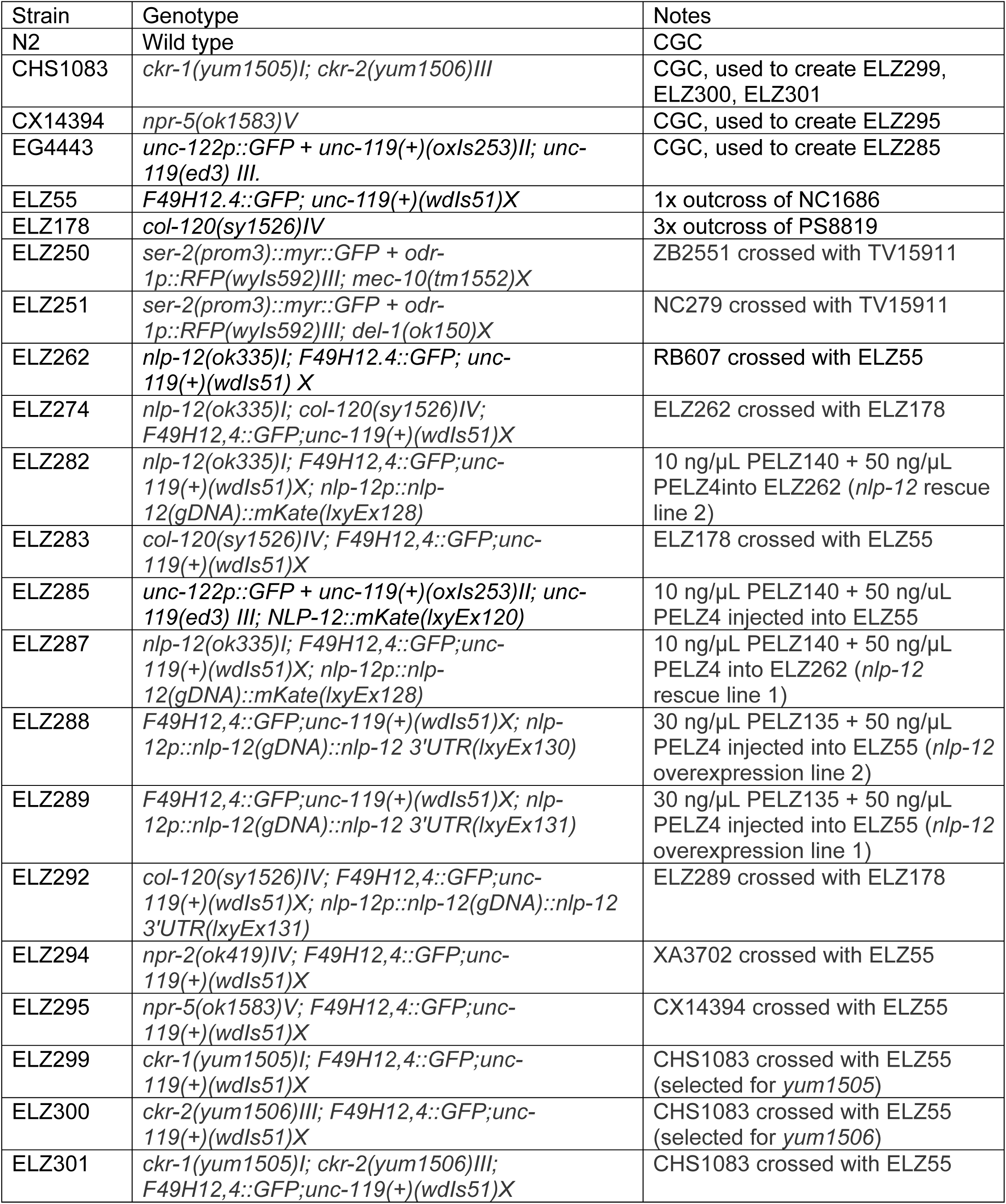
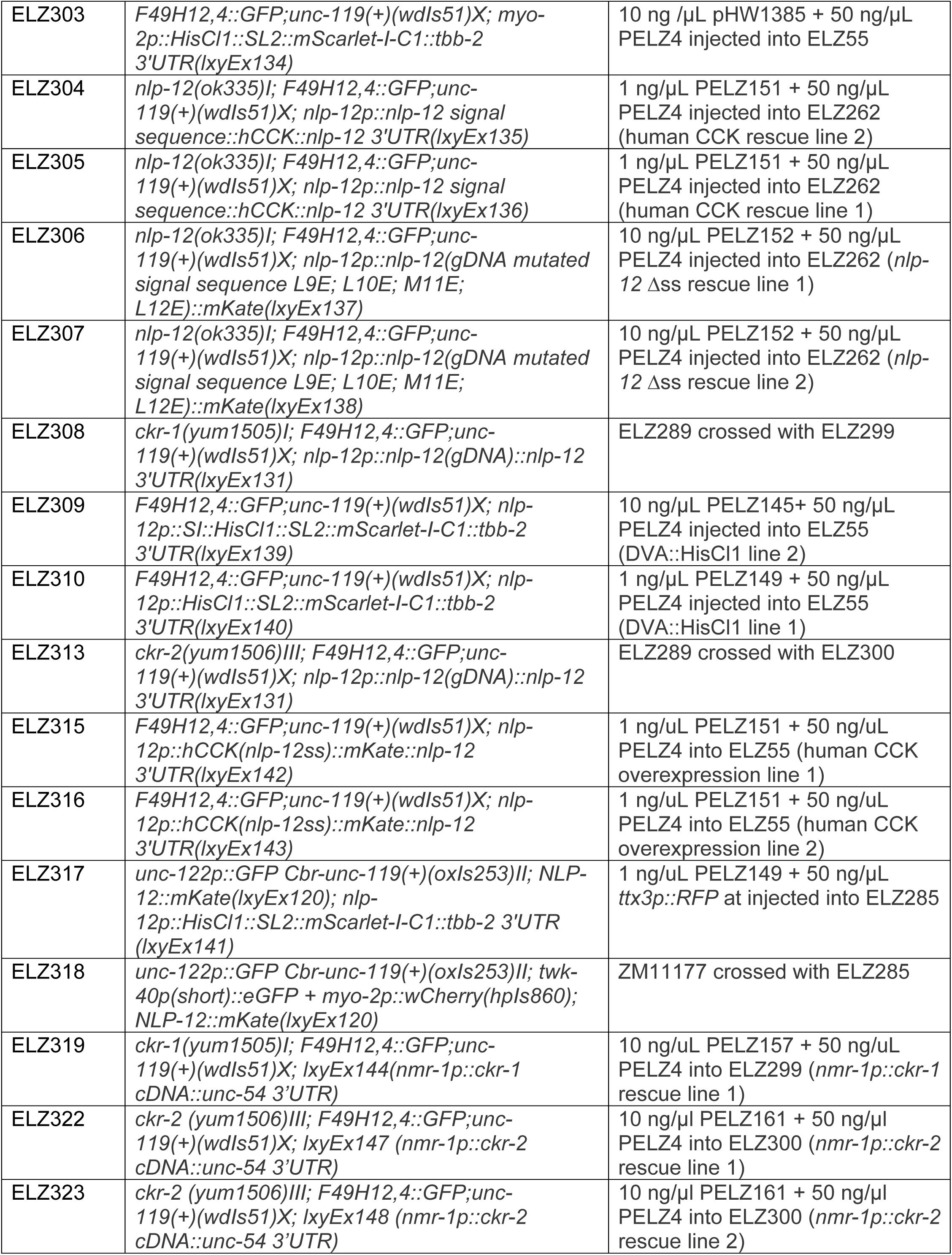
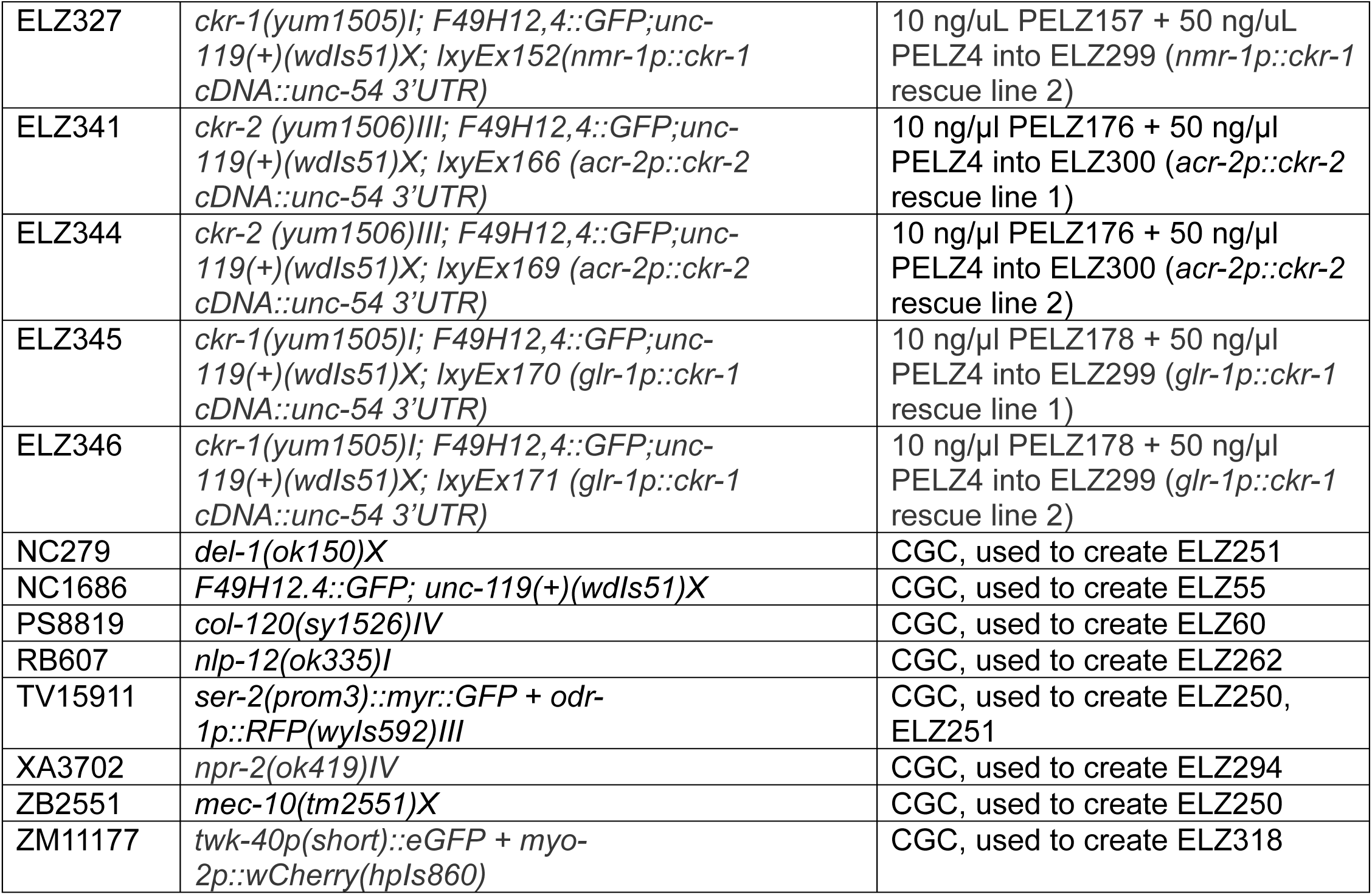
List of Strains.

**Table S2:** List of Plasmids.

| Plasmid number | Plasmid description |
| --- | --- |
| PC115339 | CCK ORF vector (Human) [Applied Biological Materials] |
| PELZ4 | *ttx-3p::GFP* |
| PELZ135 | *nlp-12p::nlp-12(gDNA)::nlp-12 3'UTR* |
| PELZ140 | *nlp-12p::nlp-12(gDNA)::mKate::nlp-12 3'UTR* |
| PELZ145 | *nlp-12p::Sl::HisCl1::SL2::mScarlet-I-C1::tbb-2 3'UTR* |
| PELZ149 | *nlp-12p::HisCl1::SL2::mScarlet-I-C1::tbb-2 3'UTR* |
| PELZ151 | *nlp-12p::nlp-12 signal sequence::hCCK::nlp-12 3'UTR* |
| PELZ152 | *nlp-12p::nlp-12(gDNA mutated signal sequence L9E; L10E; M11E; L12E)::mKate* |
| PELZ157 | *nmr-1p::ckr-1 cDNA::unc-54 3'UTR* |
| PELZ161 | *nmr-1p::ckr-2 cDNA::unc-54 3'UTR* |
| PELZ176 | *acr-2p::ckr-2 cDNA::unc-54 3'UTR* |
| PELZ178 | *glr-1p::ckr-1 cDNA::unc-54 3'UTR* |
| pHW1385 | *myo-2p::HisCl1 cDNA::SL2::mScarlet-I-C1::tbb-2 3'UTR* [Addgene] |

**Table S3:**
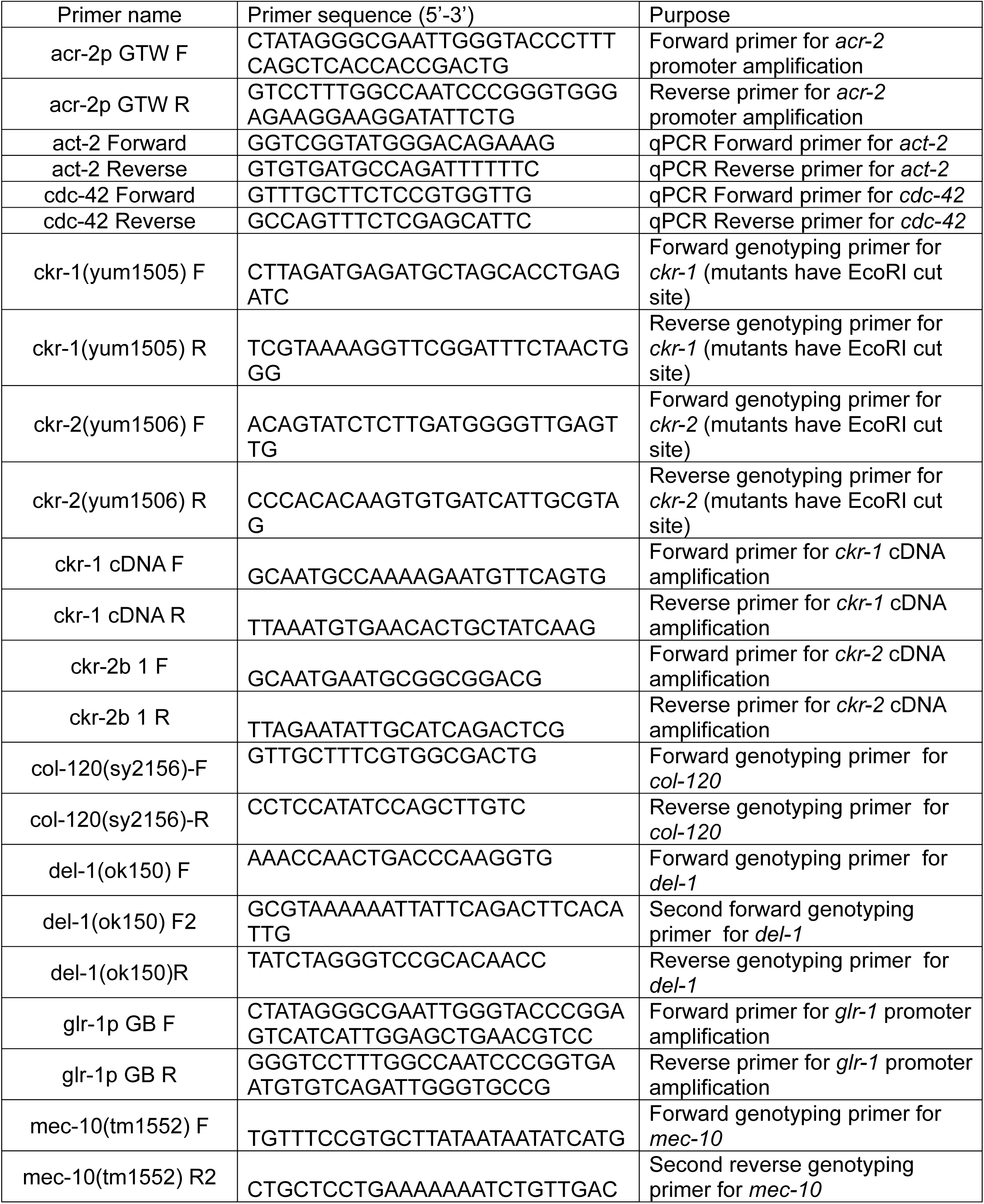
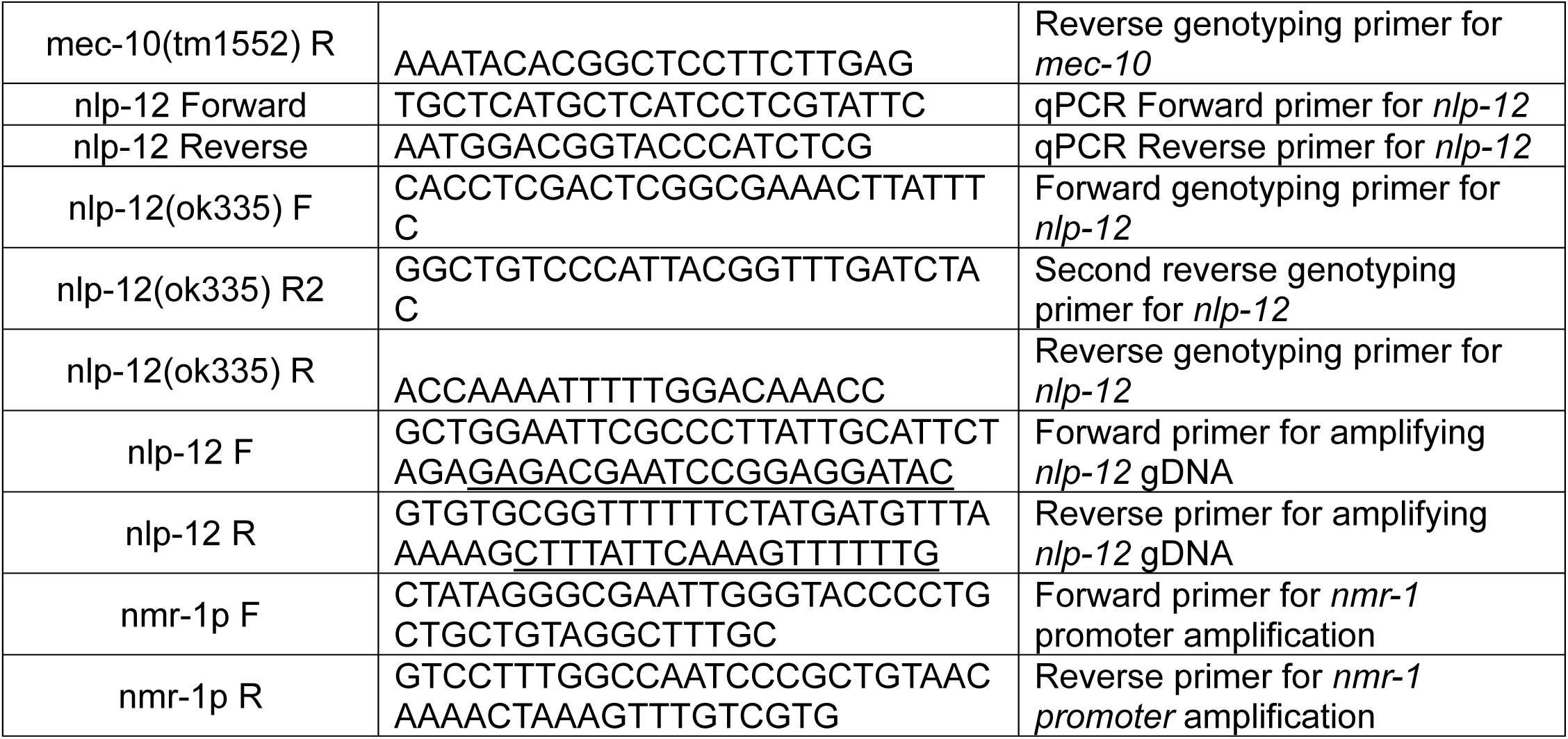
List of Primers.

## References

Bhattacharya R, Touroutine D, Barbagallo B, Climer J, Lambert CM, Clark CM, Alkema MJ, Francis MM. 2014. A conserved dopamine-cholecystokinin signaling pathway shapes context-dependent caenorhabditis elegans behavior. PLoS Genet. 10(8):e1004584.

Brockie PJ, Mellem JE, Hills T, Madsen DM, Maricq AV. 2001. The c. Elegans glutamate receptor subunit nmr-1 is required for slow nmda-activated currents that regulate reversal frequency during locomotion. Neuron. 31(4):617–630.

Callisaya ML, Blizzard L, Schmidt MD, McGinley JL, Srikanth VK. 2010. Ageing and gait variability--a population-based study of older people. Age Ageing. 39(2):191–197.

Chatzigeorgiou M, Yoo S, Watson JD, Lee WH, Spencer WC, Kindt KS, Hwang SW, Miller DM, Treinin M, Driscoll M et al. 2010. Specific roles for deg/enac and trp channels in touch and thermosensation in c. Elegans nociceptors. Nat Neurosci. 13(7):861–868.

Chemelli RM, Willie JT, Sinton CM, Elmquist JK, Scammell T, Lee C, Richardson JA, Williams SC, Xiong Y, Kisanuki Y et al. 1999. Narcolepsy in orexin knockout mice: Molecular genetics of sleep regulation. Cell. 98(4):437–451.

Chen L, Liu Y, Su P, Hung W, Li H, Wang Y, Yue Z, Ge MH, Wu ZX, Zhang Y et al. 2022. Escape steering by cholecystokinin peptidergic signaling. Cell Rep. 38(6):110330.

Chen L, Su P, Wang Y, Liu Y, Chen LM, Gao S. 2024. Ckr-1 orchestrates two motor states from a single motoneuron in c. Elegans. iScience. 27(4):109390.

Choi S, Taylor KP, Chatzigeorgiou M, Hu Z, Schafer WR, Kaplan JM. 2015. Sensory neurons arouse c. Elegans locomotion via both glutamate and neuropeptide release. PLoS Genet. 11(7):e1005359.

Cook SJ, Jarrell TA, Brittin CA, Wang Y, Bloniarz AE, Yakovlev MA, Nguyen KCQ, Tang LT, Bayer EA, Duerr JS et al. 2019. Whole-animal connectomes of both caenorhabditis elegans sexes. Nature. 571(7763):63–71.

E L, Zhou T, Koh S, Chuang M, Sharma R, Pujol N, Chisholm AD, Eroglu C, Matsunami H, Yan D. 2018. An antimicrobial peptide and its neuronal receptor regulate dendrite degeneration in aging and infection. Neuron. 97(1):125–138 e125.

El Mouridi S, AlHarbi S, Frokjaer-Jensen C. 2021. A histamine-gated channel is an efficient negative selection marker for c. Elegans transgenesis. MicroPubl Biol. 2021.

Fares H, Greenwald I. 2001. Genetic analysis of endocytosis in caenorhabditis elegans: Coelomocyte uptake defective mutants. Genetics. 159(1):133–145.

Fu H, Hardy J, Duff KE. 2018. Selective vulnerability in neurodegenerative diseases. Nat Neurosci. 21(10):1350–1358.

Garcia VJ, Daur N, Temporal S, Schulz DJ, Bucher D. 2015. Neuropeptide receptor transcript expression levels and magnitude of ionic current responses show cell type-specific differences in a small motor circuit. J Neurosci. 35(17):6786–6800.

Hays SE, Paul SM. 1982. Cck receptors and human neurological disease. Life Sci. 31(4):319–322.

Hu Z, Pym EC, Babu K, Vashlishan Murray AB, Kaplan JM. 2011. A neuropeptide-mediated stretch response links muscle contraction to changes in neurotransmitter release. Neuron. 71(1):92–102.

Hu Z, Vashlishan-Murray AB, Kaplan JM. 2015. Nlp-12 engages different unc-13 proteins to potentiate tonic and evoked release. J Neurosci. 35(3):1038–1042.

Hums I, Riedl J, Mende F, Kato S, Kaplan HS, Latham R, Sonntag M, Traunmuller L, Zimmer M. 2016. Regulation of two motor patterns enables the gradual adjustment of locomotion strategy in caenorhabditis elegans. Elife. 5.

Husson SJ, Costa WS, Wabnig S, Stirman JN, Watson JD, Spencer WC, Akerboom J, Looger LL, Treinin M, Miller DM, 3rd et al. 2012. Optogenetic analysis of a nociceptor neuron and network reveals ion channels acting downstream of primary sensors. Curr Biol. 22(9):743–752.

Jacob TC, Kaplan JM. 2003. The egl-21 carboxypeptidase e facilitates acetylcholine release at caenorhabditis elegans neuromuscular junctions. J Neurosci. 23(6):2122–2130.

Janssen T, Meelkop E, Lindemans M, Verstraelen K, Husson SJ, Temmerman L, Nachman RJ, Schoofs L. 2008. Discovery of a cholecystokinin-gastrin-like signaling system in nematodes. Endocrinology. 149(6):2826–2839.

Kampmann M. 2024. Molecular and cellular mechanisms of selective vulnerability in neurodegenerative diseases. Nat Rev Neurosci. 25(5):351–371.

Khan H, Huang X, Raj V, Wang H. 2025. A versatile site-directed gene trap strategy to manipulate gene activity and control gene expression in caenorhabditis elegans. PLoS Genet. 21(1):e1011541.

Krishna MM, Waghmare SG, Franitza AL, Maccoux EC, E L. 2025. Epidermal collagen reduction drives selective aspects of aging in sensory neurons. Aging Cell. 24(4):e14459.

Lakowski B, Hekimi S. 1998. The genetics of caloric restriction in caenorhabditis elegans. Proc Natl Acad Sci U S A. 95(22):13091–13096.

Laurent P, Ch’ng Q, Jospin M, Chen C, Lorenzo R, de Bono M. 2018. Genetic dissection of neuropeptide cell biology at high and low activity in a defined sensory neuron. Proc Natl Acad Sci U S A. 115(29):E6890–E6899.

Li W, Feng Z, Sternberg PW, Xu XZ. 2006. A c. Elegans stretch receptor neuron revealed by a mechanosensitive trp channel homologue. Nature. 440(7084):684–687.

Liang X, Dong X, Moerman DG, Shen K, Wang X. 2015. Sarcomeres pattern proprioceptive sensory dendritic endings through unc-52/perlecan in c. Elegans. Dev Cell. 33(4):388–400.

Lofberg C, Harro J, Gottfries CG, Oreland L. 1996. Cholecystokinin peptides and receptor binding in alzheimer’s disease. J Neural Transm (Vienna). 103(7):851–860.

Ludwig M, Sabatier N, Bull PM, Landgraf R, Dayanithi G, Leng G. 2002. Intracellular calcium stores regulate activity-dependent neuropeptide release from dendrites. Nature. 418(6893):85–89.

Ly K, Reid SJ, Snell RG. 2015. Rapid rna analysis of individual caenorhabditis elegans. MethodsX. 2:59–63.

Mazurek MF, Beal MF. 1991. Cholecystokinin and somatostatin in alzheimer’s disease postmortem cerebral cortex. Neurology. 41(5):716–719.

Narasimhan SD, Yen K, Bansal A, Kwon ES, Padmanabhan S, Tissenbaum HA. 2011. Pdp-1 links the tgf-beta and iis pathways to regulate longevity, development, and metabolism. PLoS Genet. 7(4):e1001377.

Nathoo AN, Moeller RA, Westlund BA, Hart AC. 2001. Identification of neuropeptide-like protein gene families in caenorhabditiselegans and other species. Proc Natl Acad Sci U S A. 98(24):14000–14005.

Peeters L, Janssen T, De Haes W, Beets I, Meelkop E, Grant W, Schoofs L. 2012. A pharmacological study of nlp-12 neuropeptide signaling in free-living and parasitic nematodes. Peptides. 34(1):82–87.

Plagman A, Hoscheidt S, McLimans KE, Klinedinst B, Pappas C, Anantharam V, Kanthasamy A, Willette AA, Alzheimer’s Disease Neuroimaging I. 2019. Cholecystokinin and alzheimer’s disease: A biomarker of metabolic function, neural integrity, and cognitive performance. Neurobiol Aging. 76:201–207.

Pokala N, Liu Q, Gordus A, Bargmann CI. 2014. Inducible and titratable silencing of caenorhabditis elegans neurons in vivo with histamine-gated chloride channels. Proc Natl Acad Sci U S A. 111(7):2770–2775.

Ramachandran S, Banerjee N, Bhattacharya R, Lemons ML, Florman J, Lambert CM, Touroutine D, Alexander K, Schoofs L, Alkema MJ et al. 2021. A conserved neuropeptide system links head and body motor circuits to enable adaptive behavior. Elife. 10.

Rossor MN, Rehfeld JF, Emson PC, Mountjoy CQ, Roth M, Iversen LL. 1981. Normal cortical concentration of cholecystokinin-like immunoreactivity with reduced choline acetyltransferase activity in senile dementia of alzheimer type. Life Sci. 29(4):405–410.

Sasidharan N, Sumakovic M, Hannemann M, Hegermann J, Liewald JF, Olendrowitz C, Koenig S, Grant BD, Rizzoli SO, Gottschalk A et al. 2012. Rab-5 and rab-10 cooperate to regulate neuropeptide release in. P Natl Acad Sci USA. 109(46):18944–18949.

Saxena S, Caroni P. 2011. Selective neuronal vulnerability in neurodegenerative diseases: From stressor thresholds to degeneration. Neuron. 71(1):35–48.

Shi A, Petrache AL, Shi J, Ali AB. 2020. Preserved calretinin interneurons in an app model of alzheimer’s disease disrupt hippocampal inhibition via upregulated p2y1 purinoreceptors. Cereb Cortex. 30(3):1272–1290.

Smith CJ, Watson JD, Spencer WC, O’Brien T, Cha B, Albeg A, Treinin M, Miller DM. 2010. Time-lapse imaging and cell-specific expression profiling reveal dynamic branching and molecular determinants of a multi-dendritic nociceptor in c. Elegans. Dev Biol. 345(1):18–33.

Stanley BG, Leibowitz SF. 1985. Neuropeptide y injected in the paraventricular hypothalamus: A powerful stimulant of feeding behavior. Proc Natl Acad Sci U S A. 82(11):3940–3943.

Taghert PH, Nitabach MN. 2012. Peptide neuromodulation in invertebrate model systems. Neuron. 76(1):82–97.

Tao L, Porto D, Li Z, Fechner S, Lee SA, Goodman MB, Xu XZS, Lu H, Shen K. 2019. Parallel processing of two mechanosensory modalities by a single neuron in c. Elegans. Dev Cell. 51(5):617–631 e613.

Tavernarakis N, Shreffler W, Wang S, Driscoll M. 1997. Unc-8, a deg/enac family member, encodes a subunit of a candidate mechanically gated channel that modulates c. Elegans locomotion. Neuron. 18(1):107–119.

Taylor SR, Santpere G, Weinreb A, Barrett A, Reilly MB, Xu C, Varol E, Oikonomou P, Glenwinkel L, McWhirter R et al. 2021. Molecular topography of an entire nervous system. Cell. 184(16):4329–4347 e4323.

Topalidou I, Cattin-Ortola J, Pappas AL, Cooper K, Merrihew GE, MacCoss MJ, Ailion M. 2016. The earp complex and its interactor eipr-1 are required for cargo sorting to dense-core vesicles. PLoS Genet. 12(5):e1006074.

van den Pol AN. 2012. Neuropeptide transmission in brain circuits. Neuron. 76(1):98–115.

Wen Q, Po MD, Hulme E, Chen S, Liu X, Kwok SW, Gershow M, Leifer AM, Butler V, Fang-Yen C et al. 2012. Proprioceptive coupling within motor neurons drives c. Elegans forward locomotion. Neuron. 76(4):750–761.

White JG, Southgate E, Thomson JN, Brenner S. 1986. The structure of the nervous system of the nematode caenorhabditis elegans. Philos Trans R Soc Lond B Biol Sci. 314(1165):1–340.

Wiesmeier IK, Dalin D, Maurer C. 2015. Elderly use proprioception rather than visual and vestibular cues for postural motor control. Front Aging Neurosci. 7:97.

Zhang N, Sui Y, Jendrichovsky P, Feng H, Shi H, Zhang X, Xu S, Sun W, Zhang H, Chen X et al. 2024. Cholecystokinin b receptor agonists alleviates anterograde amnesia in cholecystokinin-deficient and aged alzheimer’s disease mice. Alzheimers Res Ther. 16(1):109.

